# Negative sampling shapes benchmark discrimination and experimental retrieval in synthetic-lethality prediction

**DOI:** 10.64898/2026.10.25.761509

**Authors:** Zhongwei Huang, Ruohan Huang

**Affiliations:** Department of Bioengineering, University of Washington, Box 355061, Seattle, WA 98195–5061, USA; Department of Biomedical Engineering, Yale University, New Haven, CT 06520, USA

**Keywords:** synthetic lethality, gene-disjoint evaluation, negative sampling, experimental prioritization, endpoint attributes, reproducibility

## Abstract

Benchmarks for synthetic-lethality prediction compare known interactions with sampled negatives. Does performance under degree matching predict which gene pairs a model can prioritize? Across five gene-disjoint folds, matching test negatives by degree in the full labelled graph brought four methods near chance. SLMGAE states selected by benchmark test F1 had a mean area under the receiver operating characteristic curve of 0.501 on matched negatives. The same states retrieved 21 experimental hits among 198 SPIDR selections, versus 8.00 expected by chance. Comparable enrichment was not established in three other screens. Changing training negatives increased HAP1 retrieval from 3.6 to 15.8 hits for logistic regression; swapping 28 endpoint coefficients transferred most of this difference. Rewiring positive pairs while keeping the original matched negatives reduced HAP1 retrieval for logistic regression and gradient boosting. This preserved individual gene degrees but also changed their joint distribution, limiting attribution to partner identity. A density-ratio estimate closed 55–65% of the logistic-regression shortlist gap across four previously evaluated pools, with limited transfer to nonlinear models. Frozen predictions met the shortlist criterion for one of two new graph relations. Benchmark discrimination and experimental retrieval therefore warrant separate evaluation at the intended screening budget.

## Introduction

Synthetic lethality (SL) offers a route to selective cancer therapy by exploiting combinations of gene perturbations that impair viability more strongly than either perturbation alone [1]. Computational prediction can help prioritize the large space of gene pairs for experimental screening. Resources such as SynLethDB aggregate interaction evidence, and comparative benchmarks make it possible to assess models on shared inputs [2, 3]. However, recorded positives and experimentally measured candidate pools represent different sampling processes. An unrecorded pair used as a benchmark negative need not be a pair that was tested and found not to meet an experimental hit criterion.

Prior work has established several sources of misleading discrimination. In SL prediction, Seale and colleagues demonstrated gene selection bias and developed feature-based models with improved resilience [4]. Degree-aware edge sampling and bias-aware evaluation address related problems in biomedical graphs and network biology [5, 6]. Bernett and colleagues showed how network topology, protein overlap and sequence similarity affect deep protein–protein interaction predictors, including through degree-preserving controls [7]. Brière and colleagues distinguished leakage and structural shortcuts in biomedical knowledge graphs and found that cold-start evaluation did not eliminate the gap to an independent Orphanet test [8]. These studies establish the importance of controlling what a prediction task makes available to the model.

The relationship between corrected evaluation and downstream ranking is especially relevant here. Aiyappa and colleagues developed a degree-corrected link-prediction benchmark that aligned more closely with recommendation-task performance [9]. Within SL prediction, the benchmark of Feng and colleagues already included an independent test using Horlbeck’s K562 genetic-interaction screen [3]. Thus both sampling correction and experimental external evaluation have precedents. Here we ask whether a model with near-chance discrimination under a retrospective degree-matched test can still enrich for experimentally measured hits at a specified screening budget. A direct test must connect the corrected score to the retrieval behavior of the same fitted model within an explicitly measured candidate pool.

We examine this connection through three questions. First, how does test-negative composition change discrimination, and do the same fitted states retain value in measured experimental pools? Second, what carries a training-negative effect: learned feature weights, source gene frequencies, or the arrangement of source pairs? Third, can the resulting ranking changes be predicted from training negatives before fitting the alternative model? Coefficient swaps and source-positive rewiring address different parts of the second question. A negative-distribution classifier then tests the third, first on previously evaluated pools and subsequently on relations whose matched-model predictions had not yet been produced. These comparisons connect benchmark sensitivity to a manipulable learning process and its experimental consequences.

## Results

### Changing test negatives brings four methods close to chance in different directions

We used the published CV3 split, in which both endpoints of a test pair are absent from that fold’s training interaction graph [3]. For each of five folds, we retained the test genes and positives and replaced the Random negative list with a list matched to the positives’ joint endpoint-degree bins. Degree was calculated from the complete source-positive graph. This makes the comparison a retrospective, label-informed diagnostic, not an inductive procedure for selecting prospective experimental controls.

With each fitted model’s scores fixed, Random-trained SLMGAE, GCATSL and GRSMF changed from mean AUROCs of 0.782, 0.721 and 0.656 to 0.501, 0.502 and 0.492, respectively (Fig. 1). SL2MF changed in the opposite direction, from 0.376 to 0.519. Both matrix-factorization methods showed the same direction of change in all five folds. The primary SLMGAE comparison uses the independently trained run whose saved states also support the external analyses; the earlier, aggregate-only result is retained separately in Supplementary Table S2.

**Figure 1:**
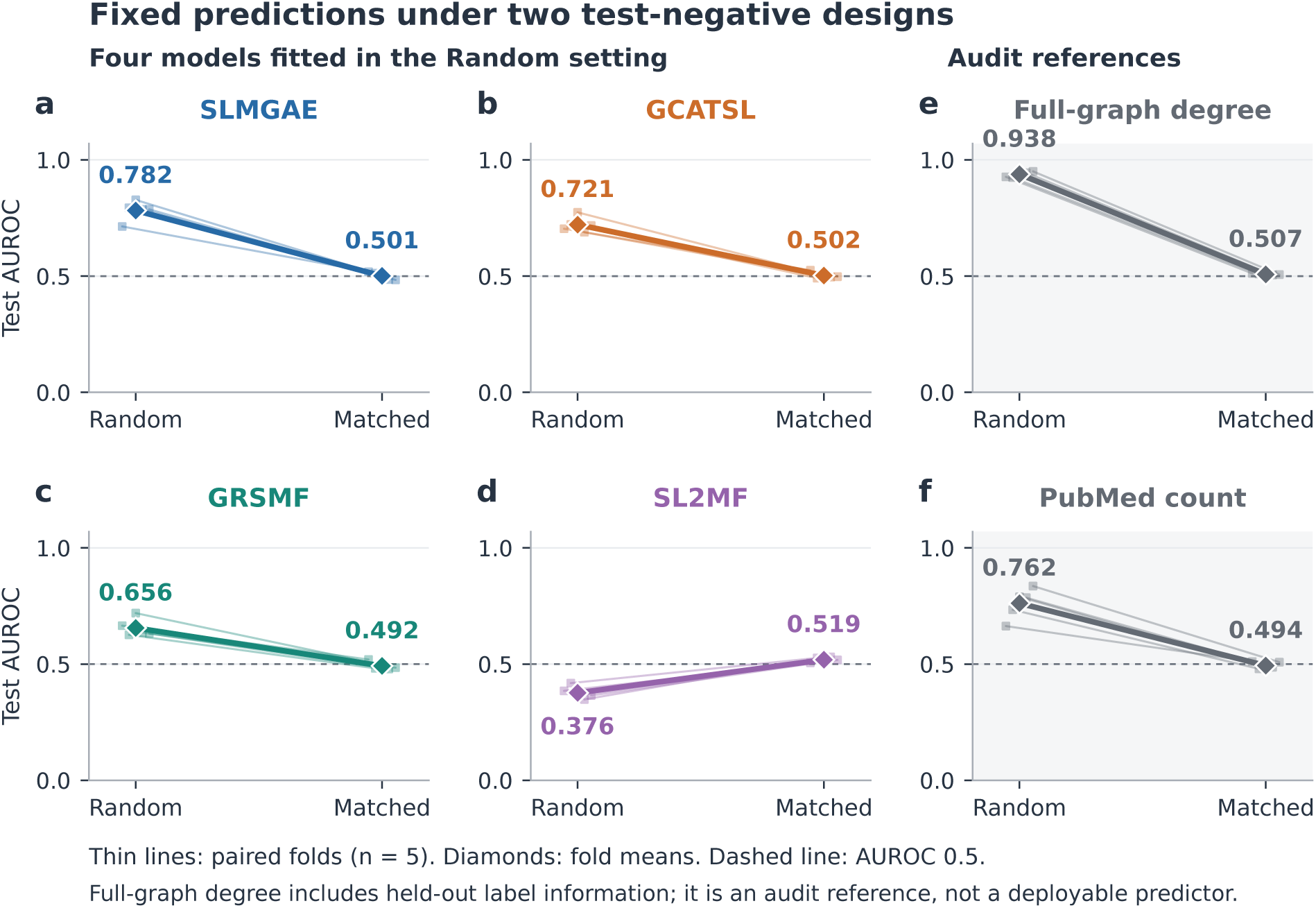
Fixed predictions under two test-negative designs. (a–d) SLMGAE, GCATSL, GRSMF and SL2MF fitted in the Random setting and evaluated with Random and degree-matched test negatives. Lines connect the same fitted state’s evaluations within a fold; diamonds and labels show unweighted five-fold means. The matched test in panels a–d uses full-source degree, including held-out positive edges, and is a retrospective diagnostic. The SLMGAE states are the same benchmark-test-F1-selected states used for external scoring. SL2MF retains its released gradient-mask convention (Methods). (e,f) Full-graph degree sum and PubMed-count sum on the same test pairs. Full-graph degree uses held-out labels and is a diagnostic reference, not an inductive predictor. Dashed lines indicate AUROC 0.5. Folds share training data and are not independent biological replicates.

SL2MF provides a counter-directional control. GRSMF scores were positively associated with full-graph degree, whereas SL2MF scores were negatively associated with it (mean Spearman correlations approximately +0.307 and −0.285). Matching the comparison pairs brought both methods closer to chance. Test composition therefore altered apparent discrimination in both directions across different model families, rather than uniformly exposing an inflated high score.

Two fixed audit references characterized the corresponding population difference. Full-graph degree sum changed from AUROC 0.938 to 0.507, and PubMed-count sum from 0.762 to 0.494. Because full-graph degree includes held-out positive labels, it serves as a label-informed diagnostic. Together, the model and reference comparisons show how strongly the negative population determines measured CV3 discrimination.

### Crossed evaluation distinguishes test composition from training degeneration

The two neural models were trained separately with Random or matched negatives, and each selected state was scored against both test lists. This 2 × 2 design separates a change in the comparison population from a change in the fitted model (Table 1). For SLMGAE, both training arms had near-chance matched-test AUROC, while Random-test AUROC remained above chance. For GCATSL, switching from Random to matched training reduced Random-test AUROC from 0.721 to 0.531. On the matched test, the two training arms scored 0.502 and 0.504, respectively.

**Table 1:** Crossed training and testing. Values are unweighted five-fold mean AUROC from the saved selected states. Each row uses identical predictions for its two test columns. Folds share training observations. Near-chance denotes proximity to 0.5 and carries no claim of statistical equivalence.

| Model | Training negatives | Random test | Matched test |
|---|---|---|---|
| SLMGAE | Random | 0.78195 | 0.50060 |
| SLMGAE | Matched | 0.71917 | 0.50240 |
| GCATSL | Random | 0.72130 | 0.50239 |
| GCATSL | Matched | 0.53097 | 0.50434 |

Three fixed-fold GCATSL runs with different initialization seeds developed constant training outputs at epochs 74–76 and remained constant through epoch 200 (training AUROC 0.5; Supplementary Fig. S1). Benchmark and external evaluations used earlier, nonconstant selected states. SLMGAE matched runs attained training AUROCs of 0.950–0.977 at termination and 0.563–0.914 at the selected checkpoints. Constant final scores were specific to GCATSL in this comparison (Supplementary Section S2).

A subsequent SLMGAE control selected epochs by F1 on a gene-disjoint inner validation set rather than benchmark-test F1. Across five folds and three seeds per fold, Random training yielded AUROC 0.766 on Random test negatives, 0.499 on the frozen matched list and 0.516 on a list matched after removing evaluated test-positive edges from the degree reference. A fixed epoch-99 analysis gave 0.767, 0.504 and 0.521. Thus the benchmark discrimination contrast persisted under both rules. This control does not establish whether the SPIDR hit count persists for alternative checkpoints (Supplementary Section S9). The alternative degree reference retained source edges linking test genes to training genes; it addresses direct use of evaluated test-positive edges, not strict training-graph-only deployment.

### Experimental candidate pools define distinct retrieval tasks

We evaluated the saved neural-model states on four measured screens: the HAP1 interaction map, a 27-cell-line compendium (GB27), SPIDR DNA-damage-response screening, and SCHEMATIC [10–13]. Both endpoints of every eligible pair belonged to the same official CV3 test fold, and the pair was scored only by that fold’s model. Pools, author-derived hit definitions and budgets were frozen for each task before its model scores were examined; these freezes occurred sequentially during the project, not before all experiments began.

Coverage varied markedly (Table 2). HAP1 retained 309,372 measurement records across 164 queries. GB27 retained 37 distinct pairs and only seven pairs positive in at least one cell line. SPIDR and SCHEMATIC retained 19,664 and 1,881 distinct pairs. These assay-defined populations differ in perturbations, biological backgrounds and candidate preselection; their non-calls are measured comparison observations. HAP1 retained 800 measurements of source-positive pairs while excluding the corresponding fold’s training pairs. The other three primary pools additionally excluded source-positive and benchmark selection-test overlaps. Interpretation therefore follows each task’s population and exclusion rule.

**Table 2:** Measured candidate pools after mapping and task-specific exclusions. Positives follow each screen’s frozen author-derived definition. Budget counts selection slots. Removing 800 HAP1 records overlapping known source positives left 377 label-A positives and a matched-minus-Random LR precision gain of 0.74 pp (95% interval 0.33–1.27; Supplementary Section S8).

| Screen | Eligible population | Positive observations | Primary budget |
|---|---|---|---|
| HAP1 | 309,372 measurements; 164 queries | 382 measurements | 1,640 |
| GB27 | 37 pairs × 27 cell lines | 82 records; 7 distinct pairs | 135 |
| SPIDR | 19,664 pairs; 449 genes | 794 pairs | 198 |
| SCHEMATIC | 1,881 pairs; 209 genes | 278 pairs | 21 |

At the prespecified top-1% budget within each SPIDR fold, Random-trained SLMGAE retrieved 21 hits in 198 selections, versus 8.00 expected under random ranking of the same measured pools (Fig. 2). The pooled hit proportion was 10.61%, corresponding to 2.62-fold the random expectation. The primary equal-fold precision gain was 6.30 percentage points; its conditional random-selection test gave within-task Holm-adjusted *p* = 0.0008 across four model/training configurations. The matched-trained SLMGAE arm retrieved eight hits, while the two GCATSL arms retrieved three and five.

**Figure 2:**
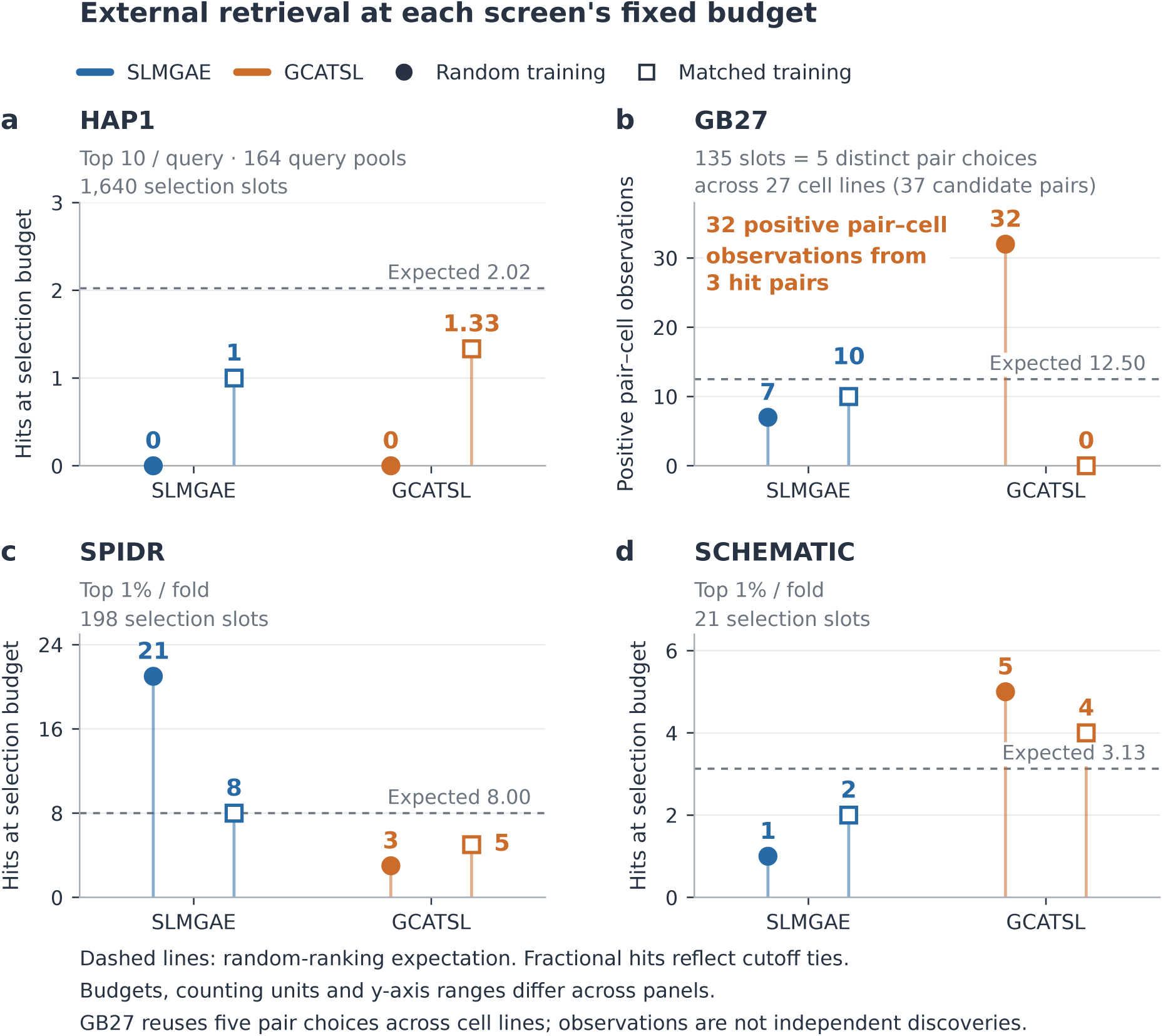
Retrieval in four measured candidate pools. Filled circles denote Random training and open squares denote matched training; dashed lines show within-pool random-ranking expectations. (a) HAP1: top 10 per query across 164 queries. (b) GB27: top one per fold–cell-line block. The 135 slots repeat five pair selections across 27 cell lines; GCATSL Random’s 32 positive pair–cell observations arise from three hit pairs. (c,d) SPIDR and SCHEMATIC: ceiling-rounded top 1% within each fold. Counts may be fractional because boundary ties are averaged. Budgets, positive prevalence and axis ranges differ; counts across panels are not a common performance scale. The figure shows descriptive counts, not confidence intervals. The 21 hits in panel c are the SLMGAE Random fold-wide top-1% result. Later SPIDR panels rank ten partners per gene using separate feature-based models.

The 21 SLMGAE hits involved 27 genes. Eight involved FZR1, but removing any single gene and reranking left the precision gain positive (3.38–6.90 percentage points). Stricter and relaxed hit thresholds, exact gene mapping and exclusion of mismatch-guide pairs retained positive retrieval contrasts (Supplementary Section S4). The enrichment thus persists across the specified gene-concentration and measurement-definition checks. These same five states had a mean matched-test AUROC of 0.501. In fold order, benchmark-test F1 selected epochs 3, 113, 59, 4 and 1; SPIDR labels did not select them. The enrichment estimate is conditional on those states and this candidate pool.

The original transported neural configurations did not establish comparable robust gains in the other screens’ primary retrieval tasks. In HAP1, Random-trained SLMGAE and GCATSL each retrieved no label-A hits in 1,640 top-10 selection slots, against 2.02 expected; matched arms retrieved one and 1.33 tie-averaged hits. In GB27, GCATSL Random produced 32 positive pair–cell observations against 12.50 expected. These observations came from three selected pair identities, and the whole-pair-profile permutation test was not significant after within-task correction (*p* = 0.3508). In SCHEMATIC, GCATSL Random retrieved five hits against 3.13 expected, with corrected *p* = 0.404; SLMGAE Random retrieved one. The small numbers of selectable pairs or positives in some tasks preclude treating lack of significant enrichment as proof of no meaningful benefit.

A separate in-domain control showed that the eligible HAP1 task contained learnable signal. LR fitted to HAP1 measurements from other genes, using 59 non-SL knowledge-graph features, retrieved 11 hits versus 2.02 expected; its precision gain was 0.547 pp (two-learner-adjusted 97.5% interval 0.140–1.009). HGB retrieved three hits. This control changes both learner and training domain, so it establishes task learnability without isolating the cause of external transfer performance (Supplementary Sections S5–S6).

### Training negatives redirect feature use and experimental retrieval

We fitted LR and histogram gradient boosting (HGB) to the same SynLethDB training positives and 59 non-SL knowledge-graph features, changing only the negative pairs. Ten redraws per design compared uniformly sampled Random negatives, full-source-degree matching and training-partition-degree matching. The latter uses only source positives within each training partition.

Training-partition matching raised LR’s mean HAP1 top-10 retrieval from 3.6 to 15.8 hits across 164 queries and 1,640 slots, versus 2.02 expected under random ranking (Fig. 3). The query-level precision-gain difference was 0.744 percentage points (pp; 95% interval 0.329–1.268). Every matched redraw retrieved 13–18 hits, exceeding every Random redraw’s 1–5. HGB gave 3.9 versus 12.3 hits, a difference of 0.512 pp (0.159–0.909). Full-source matching gave similar results. These are label-A, top-10 comparisons within the same HAP1 pool. Both feature learners used a common 59-feature representation; the transported neural models described above did not show the same gains. The comparison changes both model architecture and representation, precluding attribution to either alone.

**Figure 3:**
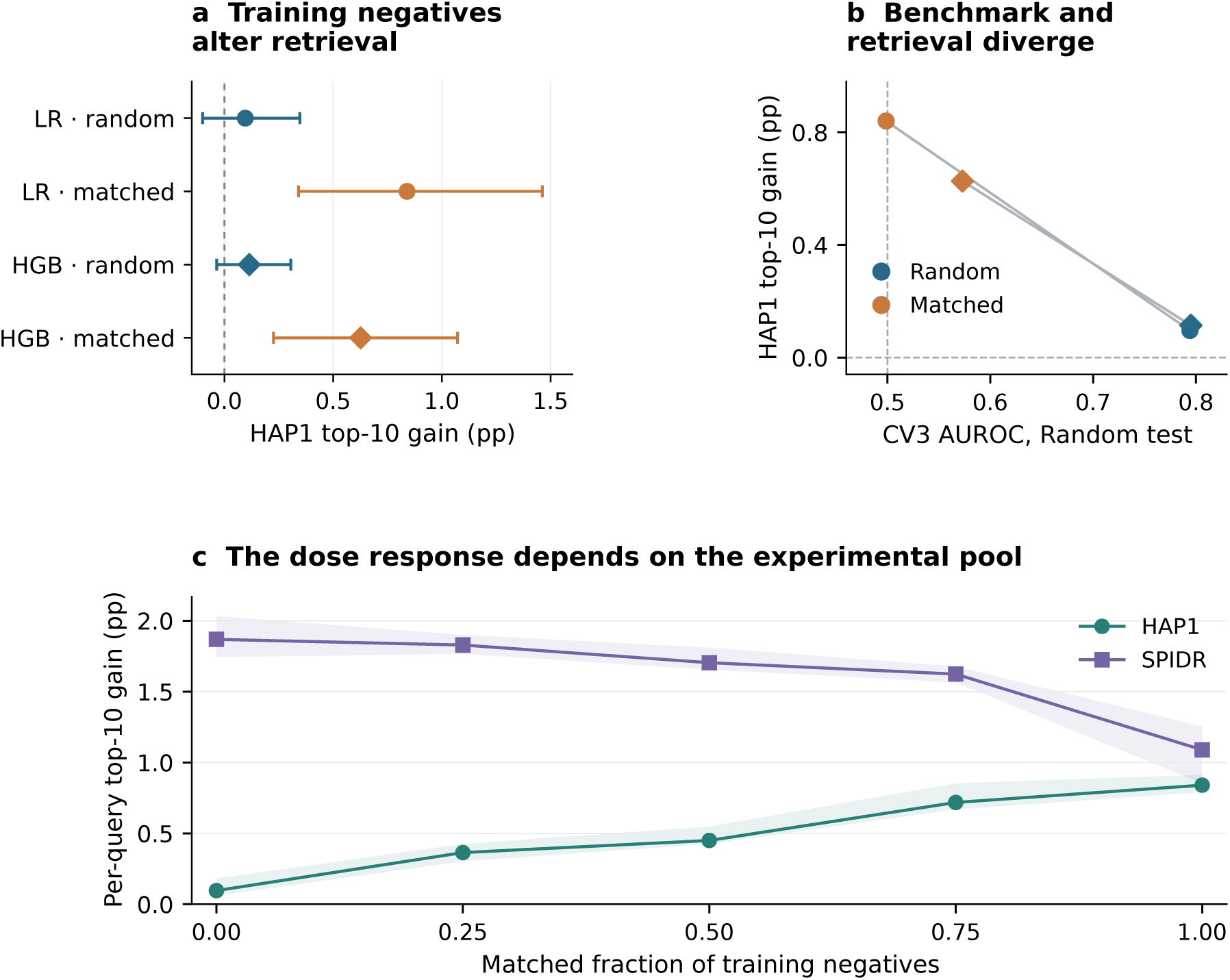
Changing training negatives redirects experimental retrieval. (a) HAP1 precision gains under two training-negative designs, with 95% query-and-redraw bootstrap intervals from ten redraws. (b) The same designs ordered by Random-test CV3 AUROC and HAP1 gain; lines join designs within each learner. Circles denote LR and diamonds HGB; blue and orange denote Random and training-degree-matched negatives. (c) LR mean-query top-10 gain across matched-negative fractions in HAP1 and SPIDR. Shading gives the range of five redraws at each dose, not biological uncertainty. SPIDR per-query retrieval differs from the fold-global top-1% task in Fig. 2. Full diagnostics are in Supplementary Section S8.

Benchmark discrimination ranked the designs opposite to HAP1 retrieval. LR’s Random-test AUROC fell from 0.794 to 0.499 as HAP1 hits increased; HGB’s fell from 0.795 to 0.573. Even HAP1 within-query AUROC was slightly higher for Random-trained LR (0.600 versus 0.571), emphasizing the distinction between whole-ranking discrimination and retrieval at a small experimental budget.

In the remaining SPIDR analyses, each of 449 genes contributes its ten highest-ranked partners (about 11% of its list, compared with about 0.5% for HAP1). This per-gene task differs from the neural fold-wide top 1% in Fig. 2. At three aligned fractional budgets, none produced intervals excluding zero in opposite directions for both screens (Supplementary Table S8). The representation contains 28 endpoint attributes—minimum and maximum degree under 14 auxiliary knowledge-graph relations—and 31 pair-relation or shared-neighbour features. Increasing the matched fraction through 0, 0.25, 0.5, 0.75 and 1 raised LR HAP1 hits through 3.6, 8.0, 9.4, 13.8 and 15.8, while reducing SPIDR per-gene precision gain from 1.87 to 1.09 pp. Removing the endpoint block from Random-trained LR likewise increased HAP1 retrieval to 12.2 hits and reduced SPIDR gain. HGB and MLP supported parts of this pattern, but neither strict dose monotonicity nor LR’s ablation asymmetry generalized to every learner (Supplementary Sections S8 and S12). At the original SPIDR fold-wide 1% budget, Random- and training-degree-matched LR instead retrieved 19.0 and 18.8 hits. The contrast at ten partners per gene therefore depends on the ranking task (Supplementary Sections S8 and S10).

### Exchanging endpoint weights transfers the LR contrast; rewiring changes HAP1 retrieval

We first intervened on fitted LR models. After mapping coefficients to a common original-feature scale, we exchanged the 28 endpoint coefficients between Random- and training-degree-matched models while retaining the 31 pair-feature coefficients. Installing matched-model endpoint coefficients in the Random model increased HAP1 hits from 3.6 to 14.1; the reverse swap reduced the matched model from 15.8 to 2.6. SPIDR per-gene retrieval moved oppositely, from 265.3 to 231.8 and from 232.8 to 272.4 hit incidences.

The symmetric factorial endpoint effect was 0.723 pp (95% interval 0.338–1.210) in HAP1, compared with 0.021 pp (−0.125–0.174) for the pair-feature block. In SPIDR, the endpoint effect was −0.814 pp (−1.395 to −0.220), with a pair-block effect of 0.090 pp (−0.171–0.339; Fig. 4). Norm-preserving swaps reduced the HAP1 endpoint effect to 0.332 pp. Thus a specified coefficient block carries most of the training-design contrast within LR; the factorial ratios are not biological mediation estimates.

**Figure 4:**
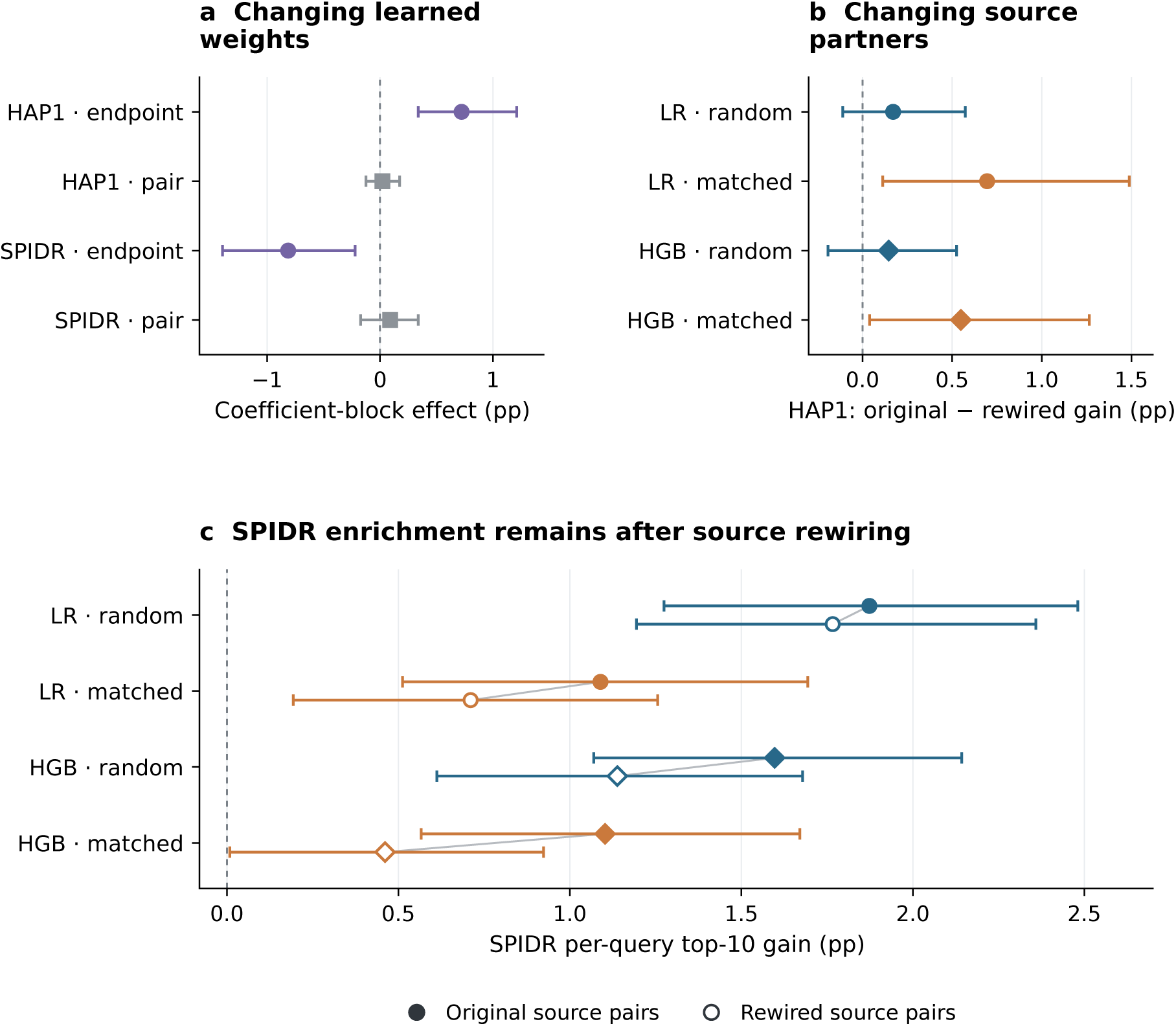
Interventions on learned weights and source pairing structure. (a) LR endpoint- and pair-coefficient effects from the symmetric swap decomposition, with 95% bootstrap intervals. (b) HAP1 gain lost after degree-preserving source-positive rewiring, with Bonferroni-adjusted intervals for eight contrasts (99.375% per contrast). Gene-level positive counts and the original training-negative lists are fixed. (c) SPIDR gains before and after rewiring, with descriptive 95% intervals; filled/open markers denote original/rewired positives. Colours denote the fixed Random or matched-negative list, and circles/diamonds denote LR/HGB. Only the two HAP1 fixed-M contrasts excluded zero after correction. Rewiring changed source partner identities and the degree-bin distribution relative to the original matched negatives. Panel c uses LR/HGB top ten per gene. Panels a and b/c use ten and five redraws, respectively. Separate gain-interval overlap is not a test of the paired contrast; full contrasts are in Supplementary Section S14.

We next changed the source relationships themselves. Degree-preserving rewiring within each training fold retained every gene’s positive count while replacing 88.9–92.6% of source edges. The original negative lists, representation and learners remained fixed. Across five folds and the first five negative redraws, HAP1 retrieval with the original matched-negative lists fell from 15.8 to 4.4 hits for LR and from 11.6 to 2.6 for HGB. The respective reductions were 0.695 and 0.549 pp; Bonferroni-adjusted bootstrap intervals across eight pool–learner–negative-list comparisons excluded zero (0.113–1.488 and 0.040–1.265 pp). Each fold and each leave-one-fold-out summary retained positive point-estimated reductions. These five-redraw comparisons are separate from the ten-redraw weight-swap series.

In SPIDR per-gene top-10 retrieval, Random-trained LR retained a precision gain of 1.766 pp after rewiring (descriptive 95% interval 1.194–2.359), compared with 1.873 pp before rewiring. This comparison did not establish equivalence. Across the eight paired contrasts, only the two HAP1 comparisons with the original matched negatives excluded zero after correction. Rewiring altered joint endpoint-degree-bin frequencies (total-variation distance 0.286–0.324 between original and rewired positives), even though it preserved each gene’s positive degree. The original matched negatives consequently need not match the rewired positives. The observed losses can reflect disruption of source pairs, loss of joint-degree alignment, or both; the experiment did not separately identify those contributions (Supplementary Section S14).

A complementary control retained the models and rearranged HAP1 target labels while preserving query and library hit margins and the measurement mask. The observed LR matched-minus-Random advantage was 12.2 hits, compared with 4.16 expected under 2,000 conditional rearrangements (*p* = 0.0005); HGB gave 8.4 versus 2.51, also *p* = 0.0005. Excluding source-positive HAP1 pairs retained the LR contrast of 0.74 pp (0.33–1.27). For both learners the observed difference exceeded every sampled null difference (minimum attainable one-sided probability 1*/*2001). Preserving these target margins alone did not reproduce the HAP1 design contrast.

The endpoint-weight effect also appeared beyond HAP1. In the Feng isogenic panel, the LR training contrast was −0.66 pp (−1.15 to −0.17), and the endpoint-block effect was −0.51 pp (−0.90 to −0.16). The observed Feng hit difference (−8.2) was close to the mean under the target-margin null (−8.89, one-sided *p* = 0.664). In HuRI protein-interaction retrieval, training matching improved LR gain by 0.291 pp (0.191–0.392), and exchanging endpoint coefficients gave an effect of 0.297 pp (0.208–0.384). A proposed degree-only rule nevertheless predicted the wrong training-benefit direction in HuRI. The functional effect transfers more broadly than that decision rule (Supplementary Sections S11–S13 and Fig. S7).

### Training-negative distributions predict part of the ranking change

Can the alternative model’s behavior be anticipated from the negative distribution itself? For equal class priors and a fixed positive distribution, the population-optimal logit satisfies 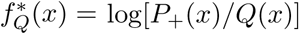. Hence

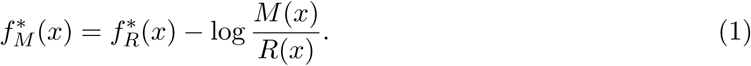

This density-ratio identity is a standard discriminative principle [22], not a new biological law. Its usefulness here depends on whether a ratio estimated in knowledge-graph feature space predicts the behavior of finite, regularized learners on genes held out from source training.

We fitted a classifier to distinguish Random from matched training negatives, without source positives, target labels or matched-model outputs. Subtracting its clipped logit from Random-trained LR scores produced a predicted matched ranking. On four previously evaluated pools, this closed 55–65% of the shortlist gap between Random and matched models: HAP1 0.592, SPIDR 0.654, Feng 0.551 and HuRI 0.571 (Fig. 5). The average closure was 0.592 and all four 95% intervals excluded zero, satisfying the prespecified aggregate shortlist criterion. HAP1 shortlist overlap rose from 0.179 for Random versus matched to 0.665 for predicted versus matched. This is a prespecified retrospective behavioral test; the reference models and pool outcomes were already known to the project.

**Figure 5:**
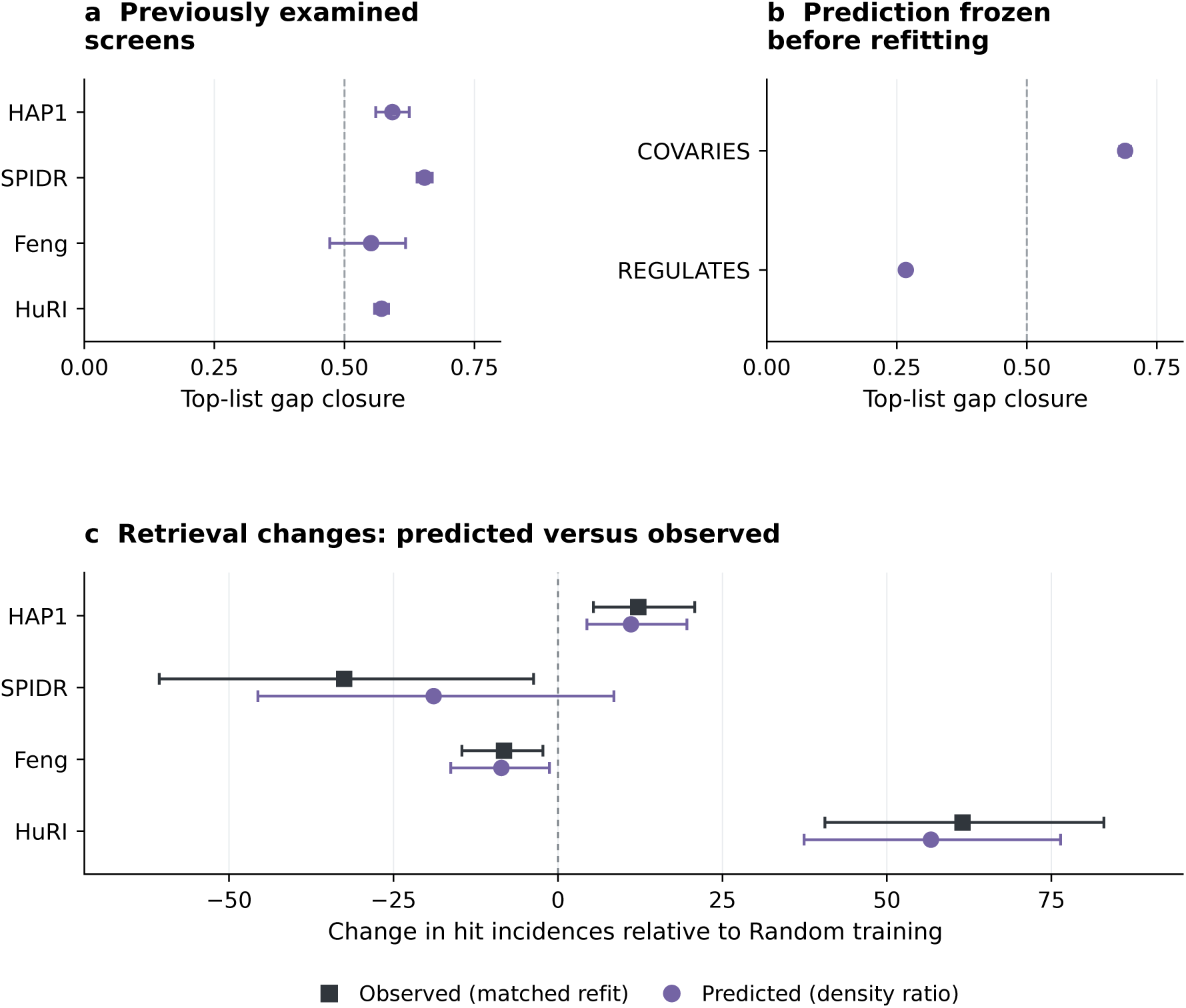
A negative-distribution model predicts part of the ranking response. (a) LR shortlist-gap closure in four previously examined pools: 1 denotes complete closure toward the retrained matched list and 0 no improvement over the Random list. (b) The same endpoint for two new relation tasks, with predictions frozen before matched-model fitting. Bars in a/b are 95% query-and-redraw bootstrap intervals; the 0.5 reference does not replace the complete task-specific criteria. (c) Observed and predicted changes in hit incidences relative to Random training, with 95% intervals. HAP1, SPIDR and HuRI use query top 10, and Feng a per-query 5% budget; counts are not comparable effect sizes across pools. The SPIDR predicted-change interval includes zero. Only COVARIES passed the frozen LR criterion in panel b; the four-pool retrieval criterion in panel c failed. Nonlinear learners and full contrasts are in Supplementary Sections S15–S16.

Retrieval prediction was less complete. The predicted hit changes were 11.1 versus 12.2 observed in HAP1, −8.6 versus −8.2 in Feng, and 56.7 versus 61.5 in HuRI. In SPIDR, the predicted change was −18.9 hits (95% interval −45.6–8.5), compared with −32.5 observed. Thus the four-pool retrieval criterion was not met, despite agreement in point-estimated directions. HGB and MLP shortlist closure was small, and a separately specified extension to SLMGAE overcorrected its rankings (Supplementary Section S15).

We then froze predictions before fitting matched models on two relation types not previously used as labels in this project. For evolutionary-rate covariation (COVARIES), LR shortlist closure was 0.689 (95% interval 0.678–0.699), satisfying the criterion of at least 0.5 with a lower bound of at least 0.4. For the second, symmetrized gene-regulation relation (REGULATES), closure was 0.267 (0.260–0.274), which did not meet the same criterion. HGB closure was approximately zero or negative. The successful result concerns prediction of a model’s shortlist, not increased experimental hit yield: COVARIES’ observed LR retrieval change was 1.0 hit (−13.9–15.7). These are computational predictions frozen before the target model existed, using two relations within the same knowledge graph, rather than prospective biological screens (Supplementary Section S16).

## Discussion

With a fixed set of model scores, changing the comparison pairs moved AUROC towards 0.5 for all four methods, from above chance for three and from below chance for SL2MF. The strongest external counterexample came from one task: the same Random-trained SLMGAE states scored 0.501 on the degree-matched benchmark and retrieved 21 SPIDR hits where eight were expected. The other three primary screens did not establish comparable neural enrichment. This contrast qualifies the use of retrospective benchmark AUROC as a guide to experimental shortlisting.

The feature-based models identify one route by which training negatives can change retrieval. Exchanging endpoint coefficients moved most of the HAP1 contrast between the two LR models. Under fixed matched negatives, both LR and HGB lost HAP1 hits after positive-edge rewiring, which preserved gene counts but changed joint degree-bin frequencies. In the SPIDR per-gene task, LR still enriched for hits after rewiring. These observations concern the features, candidate pool and selection budget used here. For an experimentalist, the top ten partners can represent about one in nine candidates in SPIDR or one in two hundred in HAP1. The aligned-budget comparisons and the Feng target-margin control preclude a general rule that the cell or assay type alone determines which training negatives to use.

The density-ratio analysis adds predictive content to this account. A classifier trained only on the negative populations anticipated much of LR’s shortlist change on held-out genes. Prediction before matched-model fitting succeeded in one new relation task and fell short in the second. This distinction matters: explaining the direction of a coefficient shift, predicting the selected candidates and choosing the training design with the best experimental yield are progressively different questions. The additive score correction tracked the difference between the two fitted LR coefficient vectors (Supplementary Section S15). Finite-sample fitting, interactions or score calibration could explain its weak performance in nonlinear learners; this study did not separate those possibilities. The SPIDR retrieval interval also crossed zero even when the LR shortlist shifted in the predicted direction.

These findings extend work on SL selection bias, degree-aware sampling, source-network controls and corrected evaluation [4–9]. Neither degree-preserving rewiring nor the population log-density-ratio identity is new. The contribution is their connection to measured retrieval through specified training and coefficient interventions, together with an empirical test of behavior prediction beyond the source-training genes. The theoretical derivations and simulations provide an interpretive model under stated assumptions; they do not supply a universal degree threshold for choosing negatives (Supplementary Section S17).

Several boundaries define this interpretation. Source rewiring preserves single-gene exposure but changes joint endpoint and pair-feature distributions. Auxiliary knowledge-graph coverage, reused pools, shared partners and related tasks limit independence; folds and negative redraws are not biological replicates. The neural models were transported to four primary SL screens using states selected by benchmark-test F1. Additional analyses of the Desjardins/Bowlan and Feng isogenic panels probed context-specific retrieval; HuRI tested transfer to protein interactions (Supplementary Sections S11–S13). Full-source degree matching used held-out labels. Later validation-selected controls retained the discrimination contrast, but do not turn full-source matching into a prospective deployment procedure. The computational prediction tasks remove direct target-relation features while retaining other information from the same graph. These limitations constrain generalization, rather than erase the within-design contrasts.

The resulting practical strategy is a linked audit: evaluate sensitivity to the comparison population, establish training viability, measure retrieval at the intended budget, and intervene on information that carries the training effect. Density-ratio prediction can supplement this audit for compatible learners, with its accuracy checked rather than assumed. A benchmark score alone cannot tell whether a model is useful for a particular screen; an external enrichment alone cannot establish which source relationships it has learned.

## Methods

### Benchmark data and degree-matched comparisons

We used the SL benchmark’s 9,845-gene universe, 35,913 unique non-self source-positive pairs and official five-fold CV3 1:1 positive-to-negative split [3]. Genes were partitioned using shuffled five-fold splitting with random state 123. Test-positive counts were 1,179, 1,139, 1,625, 1,465 and 1,673. All positive lists and gene assignments were held fixed. For matching, each pair was assigned the ordered bins of its higher- and lower-degree endpoints, using ⌊log_2_(*d* + 1)⌋ and degree *d* from the full source-positive graph. Negative pairs were sampled within the appropriate training or test node partition to match the joint-bin distribution, excluding known positives and duplicate pairs. Sparse training bins required 2/1/2/4/2 fallback selections across folds; test bins required none. Frozen pair lists, rather than a claim of byte-identical stochastic regeneration, define the comparisons.

All model inputs used the frozen negative-pair lists (Supplementary Section S7). Full-graph degree sum and the 21 October 2026 gene2pubmed count sum were evaluated on identical pairs. Training-graph degree is zero for all CV3 test genes by construction. None of the full-source diagnostics is presented as an inductive predictor.

### Models, training and saved predictions

SLMGAE, GCATSL, GRSMF and SL2MF were executed through the benchmark implementations with their released settings [15–18]. SLMGAE used learning rate 0.001, hidden dimensions 512/256, dropout 0.2 and the original maximum 300 epochs and early-stopping rule. GCATSL used 200 epochs; missing neighbourhood caches were reconstructed using the original float64 recurrence. GRSMF used 10 iterations and SL2MF a maximum of 100 with training-loss stopping. Complete settings, input hashes and implementation notes are archived with the protocols.

The neural models were fitted under both negative-training conditions. The strict improvement of primary benchmark-test maximum F1 determined each selected state; the alternate test list did not influence selection. This preserves the benchmark procedure but is test-based selection, not an independent validation design. Matrix-factorization models were fitted once per Random-training fold, without test-based checkpoint selection. SL2MF’s released gradient treats all non-positive entries as weight-one negatives; its sampled mask affects the stopping loss, so its training boundary is not equivalent to a purely sampled-loss formulation. All controls were retained regardless of the direction of the AUROC change.

The primary SLMGAE benchmark and external comparisons use identical saved states from the independent run. Historical aggregate results are reported separately. CPU metric acceleration was checked against the official implementation, without changing model equations. External score extraction used the frozen canonical index order *S*[min(*i, j*), max(*i, j*)]; benchmark extraction retained the official pair-list orientation. Reverse-orientation scores were archived because decoder matrices were not strictly symmetric. No external labels were used to choose these states or adjust model parameters.

### Gene-disjoint validation control

Within each outer fold’s 7,876 training genes, we sampled 788 validation genes without replacement (seed 20261025 + *k*, *k* = 0*, … ,* 4); the remaining 7,088 genes defined the inner training set. Training and validation positives required both endpoints in their respective sets. Cross-set positives were withheld from fitting and validation labels, but supplied the observable-degree reference for matched validation negatives. The primary validation pool combined approximately equal numbers of uniform and degree-matched negatives, with one negative per validation positive. All three initialization seeds (456, 457 and 458) shared each arm’s training pair lists.

The primary state was the first maximum of validation-pool F1 over trained epochs. Early stopping monitored improvements in validation F1 or validation-positive NDCG@10, with patience 20, at least 101 epochs and at most 300. NDCG ranked all 9,845 gene columns using only validation positives as relevant partners. Outer-test labels did not guide state selection; auxiliary GO/PPI views retained the benchmark’s full gene universe. Fixed epoch 99, the final epoch, and F1 selection on the uniform or matched validation subsets supplied sensitivity checks (Supplementary Section S9).

### External labels, mapping and overlap rules

Experimental files were mapped to the benchmark universe using the archived identifier and alias rules. Both genes had to lie in the same held-out fold. HAP1 retained screen, query, library-gene and medium identity and used label A, the authors’ released SL list, as primary; four alternative interpretations of the paper and repository definitions were retained as sensitivities [10]. Labels were not propagated to every measurement of the same unordered pair. This yields 382 positive measurements, not 413 obtained by such propagation.

GB27 used the authors’ cell-line-specific hit calls [11]. SPIDR used the frozen sensitive GEMINI score threshold ≤ −1 [12]. SCHEMATIC used membership in the authors’ Supplementary Table 4 call list, including its pooled contexts [13]. The latter three pools excluded known source-positive pairs and any benchmark selection-test pair from either arm. HAP1 was frozen earlier with fold-training exclusions only; it retained 800 source-positive measurements, including five label-A positives. Sources, attrition counts, retained definitions and exclusions are provided in Supplementary Section S1 and the release protocols.

### Retrieval metrics and uncertainty

HAP1’s primary statistic was mean query-level P@10 minus that query’s prevalence, with each of 164 query genes equally weighted. GB27 used P@1 minus prevalence per fold–cell-line block. SPIDR and SCHEMATIC used ceiling-rounded top-1% selections within each fold and an equal-fold mean of precision minus prevalence. Boundary ties contribute their expected hits under uniform tie-breaking. For a pool with *N* measured candidates, *M* positives and budget *k*, the random-ranking expectation is *kM/N*. Pooled hit counts and equal-block precision gains describe different aggregations and are not interchanged.

HAP1 uncertainty used 2,000 paired query-cluster bootstrap draws stratified by fold. GB27 used 10,000 permutations of whole 27-cell-line label profiles within folds, preserving each pair’s repeated observations. SPIDR and SCHEMATIC used 10,000 hypergeometric random selections per fold, with the same null draws shared across configurations. One-sided Monte Carlo *p* values used (1 + #{*T*_null_ ≥ *T*_observed_})*/*10,001; Holm correction covered the four model/training configurations within each task, not a project-wide family across all screens. Leave-one-gene-out and fixed-ranking node-weight perturbations were sensitivity analyses; their percentile ranges are not confidence intervals with demonstrated population coverage. AUROC and average precision were secondary external summaries. The benchmark’s trapezoidal PR area was kept distinct from average precision.

### Training diagnostic and in-domain control

The GCATSL diagnostic fixed original fold 0, Matched training and all parameters, varying graph-level seeds 101, 202 and 303. PCA used random state 456 and input fingerprints were identical. Epoch 200 and the last 20 epochs were fixed endpoints, without selecting the most favorable seed. Training scores were obtained after updating, with dropout disabled; loss followed the original dropout-enabled step.

The HAP1 control trained only on measurements with both genes among the relevant fold’s training nodes. Its 59 features described non-SL gene–annotation and gene–gene relations, excluding SL, non-SL, synthetic-rescue and HAP1 screen-derived label/fitness fields. Fixed learners were standardized logistic regression (*C* = 1) and histogram gradient boosting (200 iterations, learning rate 0.1), with no tuning. Two-learner Bonferroni-adjusted 97.5% query-bootstrap intervals assessed precision gain. This design tests in-domain learnability rather than the cause of external transfer differences; its masking chronology and secondary analyses are documented in Supplementary Section S5.

### Training-negative redraws, dose series and feature interventions

The source-trained feature models used the same 59-feature non-SL representation as the HAP1 in-domain control. The endpoint block contains 28 columns (minimum and maximum degree over 14 auxiliary relations); 31 other columns describe pair relations or neighbourhood overlap. Negative redraws kept positives, fold assignments and learners fixed. Ten draws per design compared Random, full-source-degree matched and training-partition-degree matched negatives. The mixed-design series used matched fractions 0, 0.25, 0.5, 0.75 and 1 with five redraws each. Feature ablations removed the 28 endpoint columns, then also unnormalised overlap counts, leaving 31 and 17 columns respectively. Removing explicit degree columns does not guarantee removal of all degree-correlated information.

The feature-based MLP used two ReLU hidden layers (64 and 32 units), a logistic output, Adam with initial learning rate 0.001, L2 penalty 10*^−^*^4^ and at most 200 epochs. Inputs were imputed with training medians and standardized using the corresponding training arm. Early stopping used a stratified 10% sample-level validation subset, validation accuracy and patience 10. For the frozen-model input intervention, all endpoint columns of each pair were permuted together across the entire fold-specific screen pool, while pair-feature columns remained fixed. The same permutation was shared across training arms and redraws; it was not restricted within each query (Supplementary Section S12).

HAP1 retained its frozen label-A pool and P@10 task. The later SPIDR per-gene analysis uses each of 449 genes as a query, with both directions of a candidate pair eligible in their respective rankings; its hit totals are not counts of unique discoveries. Two-stage 95% percentile intervals resampled queries within fold and negative redraw indices, using 2,000 draws with seed 20261024. Ordinary training-design contrasts resampled draw indices independently for each arm; coefficient-swap effects used shared indices across all four coefficient combinations. For the six-query isogenic panel and the Feng panel, queries were resampled across each whole panel because some folds contained only one query. These intervals condition on the experimental pools and assume exchangeability at the chosen resampling level. Dose trends used 10,000 permutations of the 25 dose-by-redraw labels; they describe the archived fixed-pool series, not independent biological replicates. The LR three-hypothesis Holm family yielded adjusted *p* ≃ 0.00030 for each trend (Supplementary Section S8).

### Coefficient swaps and conditional nulls

For logistic score 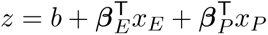, coefficients were mapped from standardized inputs to the original feature scale before block exchange. Rankings are invariant to the constant intercept within each evaluation pool. Let *G_RR_* and *G_MM_* denote precision gain with both blocks from Random or matched training, *G_MR_* the gain with matched endpoint and Random pair coefficients, and *G_RM_* the reverse. The endpoint effect is [(*G_MR_* − *G_RR_*) + (*G_MM_* − *G_RM_*)]*/*2; the pair-block effect is [(*G_RM_* − *G_RR_*) + (*G_MM_*− *G_MR_*)]*/*2. Their sum is *G_MM_* − *G_RR_* by construction. Interactions and norm-preserving swaps quantify dependence on the block context and coefficient magnitude. Neither this algebraic decomposition nor its effect ratios estimate biological mediation. Random and matched models were paired by redraw number for coefficient exchange. Their negative pairs had been sampled independently; this ordinal pairing does not imply shared negative samples. Bootstrap draws resampled these model pairs jointly across the four coefficient combinations, together with query units. Intervals were the 2.5th and 97.5th percentiles of 2,000 draws. All 100 cross-pairings of the ten Random and ten matched models provided a point-estimate sensitivity analysis for HAP1 and SPIDR.

HAP1 and Feng conditional nulls used within-fold checkerboard label swaps preserving query and partner hit margins and the observed measurement mask. For each fold with *m* positives, the chain used 20*m* attempted swaps for burn-in and 5*m* attempted swaps between stored samples; folds without admissible swaps remained fixed. HuRI used degree-preserving double-edge swaps within fold, preserving positive degree and the query set. Rankings remained fixed. Each analysis recorded 2,000 null samples and reported a one-sided exceedance probability with a plus-one correction, giving a minimum of 1*/*2001. Acceptance rates and edge-retention diagnostics are reported in the Supplement. They provide evidence of movement of the chains, not proof of independent samples or complete mixing. Null means are conditional reference values, not identified fractions of biological signal.

### Additional experimental panels and cross-task scope

The six-query isogenic panel used the released CCA rankings from Desjardins, Bowlan and colleagues [20]: FBXW7, CDK12, ARID1A, KMT2D, STK11 and KEAP1. Its primary label was each full screen’s top 50 CCA-ranked genes and its retrieval budget the smallest tested fraction meeting the prespecified expected-hit criterion (5%).

For Feng [19], we used BAGEL Bayes factors (SCORE.1) from BioGRID ORCS 2.0.18 screens 1845–1861. Query backgrounds were 1845–1856 and 1861 (TP53BP1 knockout); wild-type replicates were 1857–1860. For each gene, the mean wild-type Bayes factor was subtracted from the query-background value. Genes were ranked by descending difference across the full 18,047-gene measured intersection, breaking ties by Entrez-ID string. Top-50 membership defined the primary label; top-20 membership was a sensitivity. Mapping into the benchmark universe and restricting partners to the query’s held-out fold followed ranking, yielding 24,909 entries with 115 top-50 and 50 top-20 positives. The primary budget was ⌈0.05*n_q_*⌉ for query size *n_q_*. These rank-based endpoints differ from individually validated SL calls. Each query was scored only by its corresponding held-out fold’s source-trained model. Source-positive exclusions and alternative budgets were sensitivities.

The PPI extension trained on 7,665 mapped, non-self Lit-BM-17 interactions within the same 9,845-gene universe and five gene-disjoint folds [21]. The candidate universe was defined by the 9,243 genes also in HuRI Space III (in_space_3=1). All unordered pairs within each fold’s Space-III subset formed 8,538,813 candidate pairs, of which 3,061 were HuRI positives. Top-10 retrieval was evaluated for the 2,111 genes with at least one positive partner in their fold. Noncalled pairs served as negative reference observations; Space-III membership defines assay eligibility, not confirmed absence of an interaction for every pair.

We removed the knowledge graph’s direct interaction relation, which overlaps the PPI labels, leaving 54 features: 26 endpoint and 28 pair features across 13 auxiliary relations. Random and matched training negatives were sampled at a 1:1 ratio from the outer-training genes, excluding all mapped Lit-BM positives. Matching used ordered ⌊log_2_(*d* + 1)⌋ endpoint bins from the full or training-partition Lit-BM graph, with adjacent-bin fallback when needed. LR, HGB and MLP used the specified fixed configurations. Sensitivities excluded Lit-BM pairs from retrieval, restricted to HuRI-new interactions, varied budgets and removed GO features. Supplementary Section S13 specifies the candidate counts, feature relations, sampling seeds and interval construction.

### Degree-preserving source-positive rewiring

The source intervention used the first five archived negative redraws in each official CV3 fold. Each fold–redraw pair received an independent undirected double-edge-switch chain, with the union of its two original negative lists forbidden. We made 50|*E*| symmetric proposals, counting rejections, while preserving edge count, each node’s positive degree and the training-node partition; original edges could recur. Both negative arms used the same rewired graph and unchanged LR/HGB settings. This yielded 100 new fits. Original HGB references reused archived predictions after exact reference checks; all 50 original LR references were refitted in the same runtime as rewired LR because the historical runtime did not reproduce scores closely enough. This technical change preceded the rewired outcomes and did not change hyperparameters or endpoints.

HAP1 and SPIDR used their fixed query-top-10 pools. For each pool, learner and fixed negative list, the primary contrast was original minus rewired mean query precision gain. Ten thousand paired two-stage bootstrap draws resampled redraw indices and queries within fold, sharing samples across all comparisons. We report 95% descriptive intervals and 99.375% intervals for the eight primary contrasts (Bonferroni family level 95%). Fold and leave-one-fold-out results describe sensitivity to the partition. Pair sharing is not fully captured by query resampling. Joint feature-distribution diagnostics and chain specifications are in Supplementary Section S14.

### Density-ratio behavior prediction

For each training fold and negative redraw, a logistic discriminator assigned label zero to Random negatives and one to training-degree-matched negatives, using equal class sizes and the same non-SL feature representation. Its regularization was chosen by source-negative-only cross-validation; neither positives, target outcomes nor matched-model outputs guided this choice. Its logit estimated log[*M* (*x*)*/R*(*x*)] and was clipped at the 0.5th and 99.5th percentiles of the source-negative logit distribution. We subtracted that estimate from the Random learner’s logit to predict a matched ranking. The shortlist endpoint was the additional predicted–matched overlap over Random–matched overlap, divided by the remaining overlap gap. Overlaps were averaged across queries before forming the ratio, then averaged over redraws. The retrieval endpoint compared predicted and observed changes in hit incidences relative to Random training. Shared two-stage bootstrap samples produced 95% intervals. These analyses are retrospective with respect to the reference models and previously evaluated pools; specifications and non-LR extensions are in Supplementary Section S15.

For the subsequent computation, we used evolutionary-rate covariation and symmetrized regulation relations from the existing knowledge graph, deleted the five direct features of the target relation, and retained the same gene-disjoint folds. Every held-out gene ranked the other genes in its fold. Predictions from Random models and negative discriminators were saved and frozen before fitting any matched model on these relations. The primary COVARIES task and second REGULATES task used the same shortlist criterion; secondary retrieval used relation-held-out positives rather than new experimental measurements. Supplementary Section S16 gives the stage ordering and complete specifications. Supplementary Section S17 derives the idealized density-ratio relation and separates it from approximations for finite-sample rankings.

### Reproducibility and study scope

The suite distinguishes results recomputed from saved predictions, historical aggregate results and model diagnostics. Reproduction of 327 numerical checks and 37 complete result tables applies to the original benchmark and external analyses; the scope of the extension checks is documented separately (Supplementary Section S7).

## Data and code availability

The Bias-Aware SL Evaluation Suite v1.1.0 is archived at doi:10.5281/zenodo.23910304 [14], with code and instructions at https://github.com/booksiyideEdward/bias-aware-sl-evaluation. The suite provides predictions, benchmark pair lists, result tables and coverage manifests. HAP1 and SCHEMATIC source labels are obtained from the original sources and derived locally. Third-party files retain their source licenses; new code is MIT-licensed and original author outputs are CC BY 4.0.

Experiment-indexed tables and checks for the model interventions and transfer analyses are in additions/v3. Tables, protocols and figure-generation code for source rewiring, density-ratio predictions and simulations are in additions/v4. The extension supports checks from archived tables and predictions; rerunning all model fits additionally requires the original model inputs and feature processing. The historical SLMGAE run survives only as aggregate results (Supplementary Section S7).

## Acknowledgements and declarations

We thank the authors of the original benchmark, prediction methods and experimental screens for making their resources available. This study received no external funding. The authors declare no competing interests. The work reanalysed public data and involved no newly recruited participants or newly performed animal experiments.

Claude and OpenAI Codex assisted with software implementation, runtime troubleshooting, analysis review and manuscript drafting. The human authors are responsible for scientific claims, source verification and the manuscript’s content. AI systems are not listed as authors.

