## Supplementary Information for "Negative sampling shapes benchmark discrimination and experimental retrieval in synthetic-lethality prediction"

#### Guide to the analyses

| Main figure | Supplementary sections and archived analyses |
|---|---|
| 1 | S1–S3 and S7: fixed-score benchmark, checkpoints and seed diagnostics. S9 (E3e, E5-B): alternative degree references and independent validation selection. |
| 2 | S1, S4–S7: external tasks, hit definitions, controls and archive coverage. S10 (E1): SPIDR conditional comparisons. |
| 3 | S8 and S10 (E3b–f, E3i): negative redraws, dose, ablation and relative budgets. S11–S13: isogenic screens, feature MLP and HuRI (E7b–E10). |
| 4 | S8 (E3g, E3h): target-margin controls and coefficient exchange. S14: degree-preserving source rewiring. |
| 5 | S15 (E11): shortlist and retrieval prediction. S16 (E12a): predictions frozen before refitting. S17 (C): assumptions and simulation. |

### S1. Evaluation tasks, labels and candidate coverage

The internal task changes the negative comparison set while holding the trained state, positive pairs and CV3 test genes fixed. The five folds contain 1,179, 1,139, 1,625, 1,465 and 1,673 test positives, each paired with an equal number of Random or degree-matched negatives. These are the benchmark's released gene-disjoint folds [1]. External retrieval instead ranks measured candidates within each eligible fold or query. A pair is eligible only when both genes belong to the same fold's held-out node set; averaging predictions across other folds would not preserve this condition.

**Table S1:** Coverage of the four external tasks. Counts are pairs unless marked as measurements (m). "Strict CV3" precedes additional source-overlap exclusions.

| Task | Source pool | Mapped pool | Strict CV3 | Primary pool and budget |
|---|---|---|---|---|
| HAP1 | 3,934,515 m after QC | 1,547,513 m | 309,372 m | 309,372 m; 382 positive m; 10 per query |
| GB27 | 472; 117 ever positive | 230; 66 ever positive | 38; 7 ever positive | 37; 7 ever positive; 1 per fold and cell line |
| SPIDR | 149,787; 5,131 positive | 100,499; 4,259 positive | 19,949; 825 positive | 19,664; 794 positive; 1% per fold |
| SCHEMATIC | 11,621; 1,805 called | 10,261; 1,647 called | 2,069; 310 called | 1,881; 278 called; 1% per fold |

HAP1 retains the measurement identity (query screen, library gene), including direction and medium [2]. Label A recovers the authors' published SL list, yielding 382 positive measurements among 306,790 unordered pairs and 164 queries. Broadcasting labels across unordered pairs would incorrectly increase this count to 413. The pool excludes the corresponding fold's training pairs and endpoints, but retains 800 measurements corresponding to 782 source-positive unordered pairs. It therefore does not satisfy the additional whole-source and benchmark-selection-pair exclusions applied to the other three screens. Label-A overlap among these 800 measurements is five; this distinction is retained in all independence claims.

GB27 preserves the authors' cell-line-specific calls: the primary 37 pairs yield 999 pair–cell-line records, including 82 positive records [3]. SPIDR uses the sensitive GEMINI threshold sens.score $\leq -1$ in RPE-1 TP53-knockout CRISPRi screens; the terminal `_mis` suffix is removed only for gene mapping, while perturbation identity is retained [4]. SCHEMATIC uses the 1,805 released Supplementary Table 4 synthetic-essential calls before mapping [5]. These calls include pooled contexts and are not the simple union of seven cell-line calls. Each unordered pair is evaluated once. Across tasks, an assayed non-call is a comparison observation, not a confirmed biological negative.

### S2. Model runs and state selection

Run identity is preserved when comparing the historical and extended analyses (Table S2). The SLMGAE run used for all external analyses was trained after the historical run's reusable model assets could not be recovered. It is an independent run, not a reconstruction of historical weights. Both neural models retain the benchmark's primary-test-F1 state-selection rule; the selected state is evaluated against both negative sets. GRSMF and SL2MF were each fitted once per fold without test-based state selection [6, 7].

**Table S2:** Run-specific five-fold AUROC means. Column labels indicate training/test negatives (R, Random; M, matched). A dash denotes an unperformed condition.

| Model/run | R/R | R/M | M/R | M/M | Evidence |
|---|---|---|---|---|---|
| SLMGAE historical | .78507 | .49914 | .72846 | .50777 | Aggregate only |
| SLMGAE run 01 | .78195 | .50060 | .71917 | .50240 | Saved predictions |
| GCATSL crossed run | .7213 | .5024 | .5310 | .5043 | Saved predictions |
| GRSMF control | .6558 | .4924 | — | — | Saved predictions |
| SL2MF control | .3764 | .5189 | — | — | Saved predictions |

The new SLMGAE run is identified as `slmgae_cell_external_20261024_run01`; GCATSL uses the crossed run dated 23 October 2026. SLMGAE selected epochs are 3/113/59/4/1 (Random) and 28/63/1/9/2 (matched). GCATSL selected epochs are 32/71/59/164/18 and 23/40/38/18/37, respectively. These are the states used externally, including the early selected matched-GCATSL states. SLMGAE's final training AUROC ranges of 0.983–0.990 and 0.950–0.977 do not describe every selected state; selected matched-state training AUROC spans approximately 0.563–0.914.

The matrix-factorization control retains the released configuration: GRSMF runs 10 iterations; SL2MF runs at most 100, with training-loss stopping. Both reproduce the published primary AUROC and pass the archived non-degeneracy checks. Some SL2MF ranking metrics at 50 were not exactly reproduced, so reproduction is not asserted for every reported benchmark metric. Its upward AUROC change is retained rather than treating a decrease as necessary evidence of composition sensitivity.

### S3. GCATSL seed diagnostic

The GCATSL diagnostic fixed fold 0, matched training pairs, original numerical settings and 200 epochs while varying graph-level seeds 101, 202 and 303 [8]. Feature arrays were byte-identical, PCA used random state 456, and initial variable summaries differed. The fixed endpoints were epoch 200 and the last 20 epochs; auxiliary snapshots selected by test F1 were archived separately.

**Table S3:** Fixed-endpoint training diagnostic. Each endpoint score distribution contains only 0.5.

| Seed | Constant-output onset | Constant epochs through 200 | Final AUROC | Score variance |
|---|---|---|---|---|
| 101 | 74 | 127 | 0.5000 | 0 |
| 202 | 76 | 125 | 0.5000 | 0 |
| 303 | 75 | 126 | 0.5000 | 0 |

All 600 epoch records were complete. Training scores and AUROC came from post-update evaluation without dropout; training loss came from the original dropout-enabled step. The benchmark's maximum F1 of 0.6667 and trapezoidal PR-AUC of 0.75 at the constant endpoint reflect threshold/curve conventions, not discrimination. This diagnostic establishes repeated degeneration under the tested configuration, without identifying an optimization mechanism or replacing the earlier selected states used externally.

### S4. External sensitivity analyses and dependence

All four configurations share each task's candidate pool, labels and budget. Boundary ties contribute expected hits under uniform random tie-breaking. HAP1 first averages within query and then gives the 164 query genes equal weight; each currently has one retained screen. SPIDR and SCHEMATIC use fold-wise ceiling-rounded 1% budgets of 39/37/45/37/40 and 3/3/4/5/6. Their primary statistic averages each fold's precision minus its candidate prevalence, so it differs from the pooled hit proportion. Scores from different folds are not ranked together.

HAP1 uses 2,000 paired query-cluster bootstrap draws stratified by fold. Labels A/B/C/D/E contain 382/382/199/164/326 positive measurements; B reproduces the current author-code rule, whereas C–E implement distinct strict interpretations of the supplementary fitness conditions. Alias deduplication leaves primary precision results unchanged. Zero observed hits can produce a degenerate empirical interval and do not establish zero population retrieval probability.

GB27 permutations exchange each pair's complete 27-cell-line label profile within its fold (10,000 draws). This preserves repeated measurements, yielding GCATSL Random $p = 0.0877$ and within-task Holm $p = 0.3508$. The 32 hits arise from three pairs. Author-label sensitivity analyses include per-cell-line BH recalculation, exclusion of single-gene-depleted records, exact-symbol mapping and restoration of the one excluded overlapping pair. The ranking is unchanged across cell lines because these models have no cell-context input.

SPIDR and SCHEMATIC use 10,000 conditional random-selection draws and Holm correction across four configurations within each task. Their 1,000 fixed-ranking node-weight perturbations assign independent exponential weights to genes and product weights to edges; reported empirical ranges are sensitivity ranges, not confidence intervals with established coverage. Leave-one-gene-out reranking further assesses concentration. SPIDR SLMGAE Random retains positive gain after every single-gene removal and gives 8/198 hits at $\leq -1.5$, 48/198 at $\leq -0.5$, 20/182 under exact mapping and 10/132 after excluding mismatch-guide pairs. SCHEMATIC exact mapping retains 5/4/1/2 hits across configurations. Pair-overlap exclusions do not establish independence of all literature or side-information sources, and folds are not biological replicates.

### S5. HAP1 in-domain positive control and oracle analysis

The positive control uses HAP1 labels from other genes to test learnability of the fixed HAP1 retrieval task. Each fold trains only on measurements whose two endpoints belong to that fold's training nodes (943,882–1,071,683 measurements; 1,050–1,302 label-A positives). The test pool is unchanged. Its 59 features derive from non-SL knowledge-graph relations: 11 gene–annotation relations provide shared counts, Jaccard similarity and endpoint annotation counts; three gene–gene relations provide adjacency, common-neighbour counts, neighbour Jaccard and endpoint degrees. Counts use the preregistered log-one-plus transformation. SL, non-SL and synthetic-rescue relations, sparse excluded relations and HAP1 screen-derived fitness/call fields are omitted.

Two learners were fixed without tuning: training-standardized logistic regression ($C = 1$, lbfgs, maximum 5,000 iterations) and histogram gradient boosting (200 iterations, learning rate 0.1, 31 leaves, minimum 20 samples per leaf, no early stopping). Scores are the logistic decision function and boosting positive-class probability. The primary criterion uses label A, KG-only features and the same query-weighted P@10 minus prevalence, with 2,000 fold-stratified query bootstrap draws. Bonferroni adjustment for two learners gives 97.5% intervals. Logistic regression passes with 11 hits versus 2.02 expected and a gain of 0.547 percentage points [0.140, 1.009]; boosting gives three hits and does not pass. Expression-augmented variants and alternative labels remain secondary.

The protocol froze at 07:59:32 UTC on 24 October 2026. Its designer had not viewed GCATSL predictions, but GCATSL analysis had already completed at 05:42:04 UTC elsewhere in the project. SLMGAE external scores were generated later, at 08:12:19 UTC. This is designer-level masking for GCATSL, not project-wide prospective registration. The comparison changes both training domain and learner, so it does not isolate the cause of transfer performance.

The prespecified source-label oracle yields five HAP1 positives among 800 source-positive measurements: risk ratio 5.12, 95% interval [0, 14.98]. Failure of its binary concordance criterion is inconclusive. A subsequent whole-mapped-pool exploration finds 25 positives among 3,911 source-positive measurements, risk ratio 5.48 [1.16, 14.38]. This exploratory extension is not a gene-held-out generalization result; absent source provenance also prevents excluding historical HAP1 contributions to SynLethDB.

### S6. Archived exploratory aggregates

The analyses in Table S4 were added after external outcomes were known. Suite v1.0.0 retains them as archived aggregates rather than re-executing them through its prediction-backed analysis workflow. They are reported as exploratory context; the manuscript's central conclusions do not depend on these comparisons or on the model-based power estimates.

### S7. Reproducibility coverage and implementation boundaries

The Bias-Aware SL Evaluation Suite v1.0.0 distinguishes prediction-backed results, archived aggregates and unexecuted analyses [9]. Saved predictions support fixed-score benchmark comparisons for the independent SLMGAE run, GCATSL and both matrix-factorization controls. The four primary external tasks use saved SLMGAE and GCATSL predictions. Release checks reproduced 327 values and 37 complete result tables within their stated tolerances. HAP1 and SCHEMATIC

**Table S4:** Post hoc aggregate records and their scope.

| Record | Archived value | Interpretation boundary |
|---|---|---|
| SPIDR richness | 7.69 expected hits; model 21; random 8 | Same 198-selection budget; score–richness Spearman mean .528; reported $p$ values compare with random, not model versus richness |
| Power at 2.63-fold enrichment | HAP1 .284; SPIDR .932; SCHEMATIC .645 | Uniform-enrichment assumption, $\alpha$ = .0125; shared-gene dependence not fully represented |
| SLMGAE Random enrichment intervals | HAP1 0 [0, 1.82]; SPIDR 2.62 [1.65, 3.74]; SCHEMATIC .32 [.01, 1.48] | Model-based archived intervals; no confirmatory cross-screen comparison |
| GB27 effective decisions | 37 pairs, 7 ever positive; 5 selected pair identities | 135 fold–cell blocks repeat five choices; power was not modelled |

source labels are downloaded and locally derived with pinned checksums rather than redistributed. The historical SLMGAE run remains aggregate-only; GCATSL seed diagnostics and the in-domain positive control are archived evidence, not suite reruns. Matrix-factorization external scores were exported but never evaluated or released. Version 1.1.0 adds archived tables and supporting materials for Supplementary Sections S8–S17, including checks of coefficient swaps, a grouped bootstrap for the source-rewiring contrasts and code for rebuilding main Figures 3–5. These checks operate on archived results; rebuilding all fits requires separately prepared features, indices and model inputs (doi:10.5281/zenodo.23910304).

Three implementation details constrain interpretation. First, neural state selection used benchmark test F1. Second, full-source degree matching uses held-out positive labels, making it a retrospective diagnostic; the full-graph degree reference is likewise label-informed. The frozen matched file originally replaced pair lists while retaining Random sparse negative-graph slots: GCATSL training rebuilt those graphs from the lists, while SLMGAE, GRSMF and SL2MF read the lists. The release corrects graph slots without changing pairs. Third, the released SL2MF gradient treats every non-positive matrix entry as a weight-one negative; sampled training negatives affect its stopping loss rather than defining the complete gradient mask. Its control therefore addresses test composition under that implementation.

Benchmark scores retain official pair orientation; external scores use the prespecified canonical order $S[\min(i,j), \max(i,j)]$, without symmetrization. The saved decoders are not strictly symmetric. Benchmark trapezoidal PR-AUC and external average precision are kept distinct throughout.

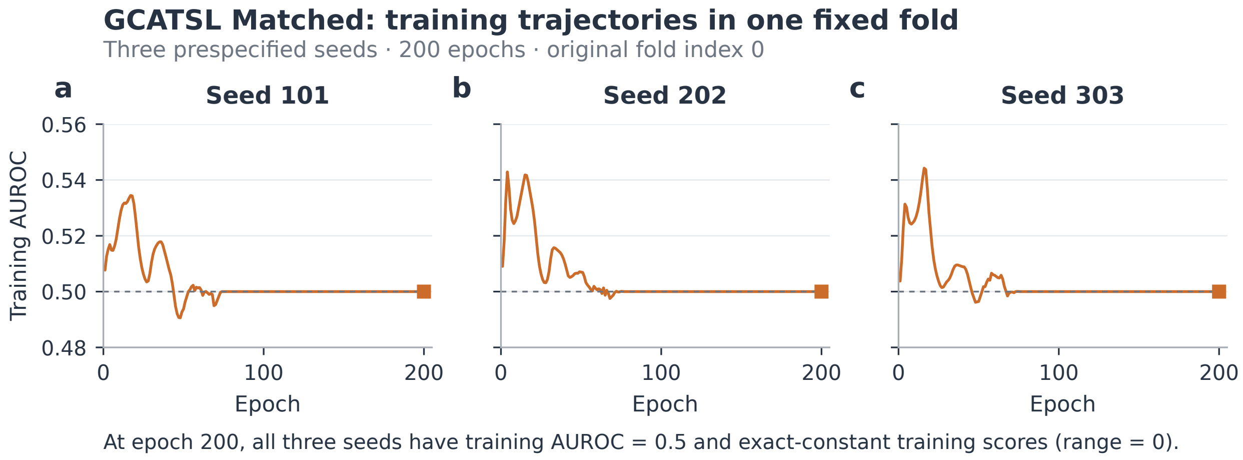


**Figure S1: Fixed-fold GCATSL training diagnostic.** All 200 training AUROCs for seeds 101, 202 and 303 under Matched training in original fold 0, without smoothing. Squares mark the fixed epoch-200 endpoint. All final score distributions are exactly constant, with variance and range zero and AUROC 0.5. These runs vary initialization within one split; they are not biological replicates. Earlier selected states, rather than the fixed endpoints shown here, were used in the external analysis.

### S8. Training-negative design and endpoint-coefficient interventions

The result tables are indexed by the archived experiment identifiers E3, E3b–E3i, E5, E7b, E8, E9 and E10. These identifiers locate source files; they do not denote independent biological replications. The source-trained feature analyses retain the original CV3 node partitions, source-positive training pairs, and 59 non-SL features. LR uses standardization and $C = 1$; HGB uses fixed hyperparameters. The exact matching recipe was checked in the original report against the frozen pair lists before generating new negative draws. Ten negative redraws were used for each of Random, full-source-degree matching and training-partition-degree matching. Seeds were 1001–1010, 2001–2010 and 3001–3010, respectively. Models from these three designs were evaluated on the same experimental pools.

**Source training and negative redraws.** The source comparison preceding the redraw experiment held the feature learner fixed. HAP1-trained LR retrieved 11 hits, source-trained LR with Random negatives three, and source-trained LR with one matched-negative draw 15. The primary HAP1-minus-source-Random contrast was 0.488 pp (95% interval 0.183–0.915). The matched arm was secondary, motivating the subsequent redraw analysis rather than establishing a general source-domain limitation.

**Table S5:** HAP1 retrieval across ten negative redraws per design. Gain is mean query precision minus query prevalence, in percentage points. Reported 95% percentile intervals resample queries within fold and negative draws (2,000 resamples). Hit ranges are redraw ranges, not confidence intervals. Random-ranking expectation is 2.023 hits.

| Learner | Training negatives | Hits, mean (range) | Gain | 95% interval |
|---|---|---|---|---|
| LR | Random | 3.6 (1–5) | 0.096 | $[-0.100, 0.347]$ |
| LR | Full-source matched | 15.9 (14–17) | 0.846 | $[0.354, 1.466]$ |
| LR | Training-degree matched | 15.8 (13–18) | 0.840 | $[0.341, 1.462]$ |
| HGB | Random | 3.9 (2–5) | 0.114 | $[-0.035, 0.305]$ |
| HGB | Full-source matched | 11.9 (8–15) | 0.602 | $[0.206, 1.063]$ |
| HGB | Training-degree matched | 12.3 (9–15) | 0.627 | $[0.226, 1.073]$ |

The LR training-degree-matched-minus-Random contrast was 0.744 pp [0.329,1.268]; HGB gave 0.512 [0.159,0.909]. All 100 cross-pairings of the ten Random and ten matched draws favored matching for HAP1. These 100 comparisons reuse 20 draws and are not 100 independent tests. The labels and pools had already been inspected in the preceding experiment. This follow-up confirms robustness to negative redraws within that screen, not independent prospective validation.

Other tasks limit a universal-benefit claim. LR SPIDR pool-level top-1% retrieval was 19.0, 19.2 and 18.8 hits for Random, full-source matched and training-degree matched, respectively (8.00 expected). GB27 P@1 totals were 57, 21 and 21 repeated pair–cell-line observations; only two changed pair choices generated the LR difference. SCHEMATIC results were unresolved. These pool-level tasks differ from the later SPIDR per-gene analysis.

**Dose series and feature ablation.** Five negative-draw replicates at each matched fraction (0, 0.25, 0.5, 0.75 and 1) yielded 25 dose–draw points per learner. LR HAP1 hit means were 3.6, 8.0, 9.4, 13.8 and 15.8; SPIDR per-gene precision gains were 1.869, 1.831, 1.702, 1.617 and 1.089 pp. The coupling summary averaged within-query score–degree correlations over HAP1, SPIDR and SCHEMATIC. LR's mean declined from 0.753 to −0.032. HGB's corresponding HAP1 means did not increase at every step: retrieval peaked at fraction 0.75 before falling at 1. The reported rank-trend tests quantify overall association rather than strict monotonicity of every learner.

The three LR one-sided dose-permutation probabilities are each 1/10001. Holm adjustment over those three hypotheses gives $3/10001 \simeq 0.000300$ for each. HGB's recorded Holm values are 0.000300 (coupling), 0.000400 (HAP1 gain) and 0.005300 (SPIDR gain). These probabilities condition on the archived dose series; the 25 points reuse data and cannot be treated as 25 independent experiments. The label-free coupling response was visible in the pre-freeze dry run, as disclosed in the source report.

A1 drops 28 explicit endpoint-degree columns; A2 also drops unnormalised overlap counts, leaving 17 columns. With Random negatives, LR A1 and A2 increased HAP1 hits by 8.6 and 5.6 and changed SPIDR gain by −0.49 and −0.83 pp. With matched negatives, HAP1 changed by only +0.4 and −0.4 hits. HGB A1 increased HAP1 hits by 1.6 even though measured coupling increased slightly, so the scalar coupling statistic is not a sufficient mediator. Remaining features can retain degree-correlated information; the interventions target the defined blocks rather than eliminating all endpoint information.

**Coefficient swap estimands.** Endpoint and pair coefficients are brought to the same original-feature scale before exchange. The four scores use Random/Random, matched/Random, Random/matched and matched/matched coefficient origins for the endpoint/pair blocks. For the primary ten matched draw pairs, mean HAP1 hits are 3.6, 14.1, 2.6 and 15.8; corresponding SPIDR per-gene hit incidences are 265.3, 231.8, 272.4 and 232.8. The endpoint effect is the average of changing that block in the two pair-block backgrounds. The pair effect is defined symmetrically. Their sum equals the total difference by algebra; the observed four scores, rather than that identity, supply the functional evidence.

For E3h, each model uses its own training medians and StandardScaler fitted to the source positives and that arm's negatives. Coefficients are converted to the original feature scale as $\beta_j = w_j/\sigma_j$ before exchange; intercepts are omitted because they do not change within-query ranking. The 28 endpoint columns are log-transformed minimum and maximum degrees for 14 non-SL KG relations, with the remaining 31 columns forming the pair block. Random draw $r$ is paired with matched draw $r$ by index; their negative sets were independently sampled, so index pairing does not imply shared sampling randomness. The original-scale score reconstruction agrees with the fitted LR decision function up to a constant (maximum absolute discrepancy $2.03 \times 10^{-6}$).

Intervals use 2,000 two-stage bootstrap replicates (seed 20261024). First, the 164 HAP1 queries or 449 SPIDR genes are resampled with replacement within their CV3 folds. Second, the ten indexed model pairs are resampled with replacement, using the same draw indices for endpoint, pair, total and interaction effects. Each replicate averages over sampled model-pair indices and then sampled query/gene units; the reported limits are the 2.5th and 97.5th percentiles. SPIDR units share pairs, so these intervals do not resolve dependence between the two endpoints' lists. Ratios of block effects to the total are point estimates without separate intervals.

The HAP1 endpoint effect is 97.1% of the total under this symmetric decomposition. A single transplant into the Random model recovers 10.5 of the 12.2 extra hits (86.1%); these are distinct estimands. The SPIDR ratio exceeds 100% because the pair-block effect partly offsets the endpoint effect. Using all 100 cross-pairings of Random and matched models gave endpoint effects of 0.73 pp in HAP1 and −0.85 pp in SPIDR. Preserving the recipient coefficient-block norm reduced the HAP1 endpoint effect to 0.332 pp, while SPIDR developed a positive interaction of 1.047 pp [0.423,1.644].

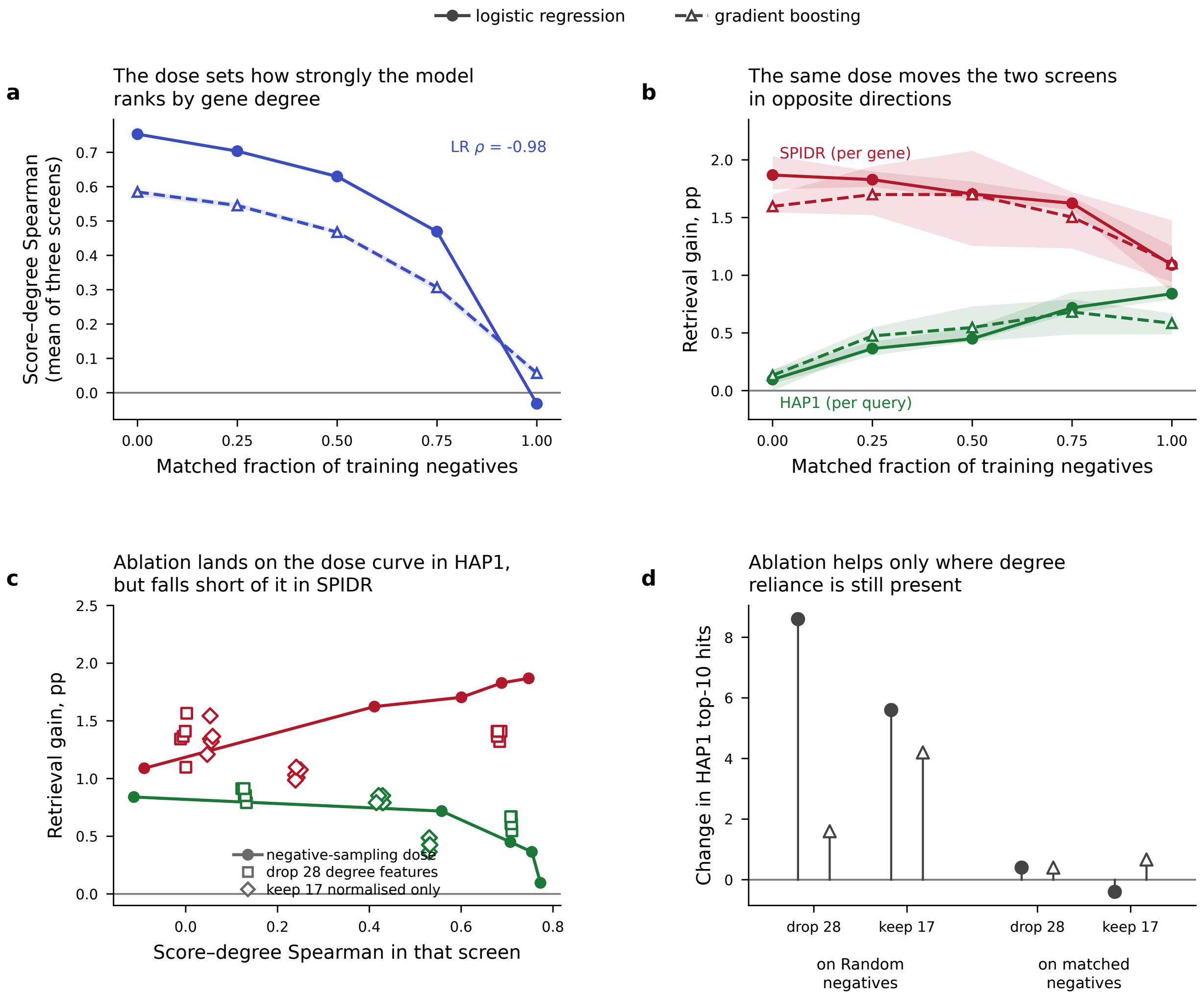


**Figure S2: Negative-sampling dose series and feature ablations.** Means over five redraws and redraw ranges for LR (solid) and HGB (dashed). (a) Score–degree coupling averaged over HAP1, SPIDR and SCHEMATIC. (b) HAP1 query and SPIDR per-gene precision gains. (c) Gain versus measured coupling; ablations are separate feature interventions. (d) HAP1 hit-count changes after removing explicit endpoint-degree columns (A1) or additionally unnormalised overlap counts (A2). Ranges summarize redraw variation, not biological uncertainty.

**Table S6:** Primary LR coefficient-swap effects in percentage points, with reported 95% intervals. Ratios to the total contrast are algebraic attributions, not causal mediation estimates.

| Effect | HAP1 | SPIDR per gene |
|---|---|---|
| Endpoint block | 0.723 [0.338, 1.210] | −0.814 [−1.395, −0.220] |
| Pair-feature block | 0.021 [−0.125, 0.174] | 0.090 [−0.171, 0.339] |
| Total contrast | 0.744 [0.329, 1.268] | −0.724 [−1.350, −0.082] |
| Interaction | 0.165 [−0.110, 0.476] | −0.136 [−0.570, 0.276] |

Thus magnitude and the other coefficient block contribute to the functional effect.

An importance-reweighting arm was cancelled before label-based assessment because its negative-sample effective size was 0.2–0.7% of the original and maximum weights were 555–3343; some target degree bins had no Random-negative support. It is not an evaluated alternative training method. The diagnostic table is retained as `E3H_reweighting_ess.csv`.

**Conditional-null and overlap controls.** HAP1 checkerboard swaps preserve query hit counts, library-entry hit counts, the observed mask and direction, with scores fixed. The five fold chains had acceptance rates 0.708–0.867 and mean retained-positive fractions 0.098–0.218. Those diagnostics do not establish full Markov-chain mixing. In 2,000 saved null samples, LR's matched-minus-Random advantage was 4.162 hits on average (SD 1.648), versus 12.2 observed; HGB's was 2.512 (SD 1.186), versus 8.4. Both reported one-sided probabilities were 1/2001. LR Random retrieval was also above this conditional null (3.6 versus 1.343, $p = 0.00850$), despite its interval against uniform random ranking including zero. Different null questions explain these results.

Removing 800 source-positive measurements, including five label-A positives, preserved the positive HAP1 contrast: LR 0.74 pp [0.33,1.27] and HGB 0.50 [0.15,0.90]. This rules out direct recovery of those retained source-positive pairs as the main explanation of the gain. It does not remove shared-gene or auxiliary-feature dependence. E3b, E3d, E3g and E3h are linked analyses of the same models and pools, not four independent validations.

**Available source tables.** The accompanying directory contains the unmodified E3B primary contrasts and redraw metrics; E3D dose, ablation and trend tables; E3G null tests, chain diagnostics and stored null samples; and E3H swap, factorial, cross-pairing and support-diagnostic tables. Source-report excerpts are retained alongside them. The corresponding tables are archived under `additions/v3` in suite v1.1.0 (doi:10.5281/zenodo.23910304).

### S9. Test-negative construction and checkpoint selection

The matched-test comparison was assessed with two controls: excluding evaluated positives from the degree information used to construct test negatives (E3e), and selecting SLMGAE checkpoints without outer-test performance (E5-B).

In E3e, the test positives were unchanged and archived score matrices were evaluated against ten redraws of each new negative construction. Let T0 denote the released Random negatives, T1 the frozen full-graph matched negatives, T2 negatives matched using only training-partition degree, and T3 negatives matched using degree over all source SL edges except the fold's held-out test positives. For Random-trained SLMGAE, mean AUROC was 0.7820 on T0, 0.5006 on T1, and 0.5169 on T3. The T3–T1 difference was 0.0163, with reported paired 95% interval [0.0110, 0.0216]; T3 means across negative redraws ranged from 0.5127 to 0.5195. Matched-trained SLMGAE gave 0.7192 on T0 and 0.5151 on T3.

T2 did not reproduce the collapse: its AUROC was 0.7795. Under CV3, training positives require both endpoints to be training genes, so all 1,969 test genes in each fold have zero training-graph degree. T2 therefore reduces to uniform sampling. T3 avoids the evaluated positive edges but uses information outside the strict training partition: approximately 10,964 test–training source edges per fold, while excluding 1,179–1,673 held-out test–test positives. Thus, T3 is a diagnostic that excludes the evaluated positives, rather than a claim that strictly training-only degrees can support nondegenerate matching for unseen genes. Its construction can still include unlabelled true interactions among the sampled negatives. E3e applies to SLMGAE; other benchmark models were not rescored against new pairs in this analysis.

**Gene-isolated validation and checkpoint selection.** E5-B retained the outer five-fold node split, constructed with `KFold(5, shuffle=True, random_state=123)` over 9,845 genes. In outer fold $k$ (zero-indexed), 788 of the 7,876 outer-training genes were sampled without replacement using seed $20261025 + k$ as validation genes, leaving 7,088 inner-training genes; the 1,969 outer-test genes were disjoint from both. Source-positive pairs with both endpoints in the inner-training set were used for fitting, and pairs with both endpoints in the validation set were used for validation. Cross-set validation–inner-training positives were excluded from fitting and validation, and used only to calculate validation genes' observable degree. Inner-training positive counts across folds were 18,703, 19,314, 18,521, 18,858 and 16,838; validation-positive counts were 310, 233, 175, 178 and 285. The corresponding excluded cross-set counts were 4,757, 4,568, 3,590, 3,718 and 4,772.

Each training arm used one negative set per fold at a 1:1 ratio to inner-training positives, shared across the three initialization seeds. Random negatives were sampled among inner-training genes; matched negatives used degree in the inner-training positive graph. The sampler bins the larger and smaller endpoint degrees by $\lfloor \log_2(d+1) \rfloor$, samples within the positive pairs' bin cells, rejects known source positives, self-pairs and duplicates, and fills exhausted cells from neighboring cells in increasing Chebyshev radius. It allows up to 60 times the required cell count in sampling attempts before expanding through radii 1–11. Random and matched training-set seeds were $20261025 + 100 + k$ and $20261025 + 200 + k$.

Validation negatives were constructed only among the 788 validation genes. Each of $V_{\mathrm{uni}}$ and $V_{\mathrm{match}}$ contained as many pairs as the validation-positive set; the latter matched degree calculated from validation–inner-training source edges, excluding the evaluated validation-positive edges. Seeds were $20261025+300+k$ and $20261025+400+k$. The shared primary validation pool $V_{\mathrm{mix}}$ combined $\lfloor n/2 \rfloor$ uniformly sampled negatives and $\lceil n/2 \rceil$ matched negatives, with duplicate pairs skipped in the matched half; the two permutation orders used seed $20261025 + 500 + k$. The benchmark training routine's test input slots contained copies of these validation lists, so no outer-test pair labels entered its metric calculation or checkpoint selection. The GO-BP, GO-CC and PPI auxiliary views still covered all 9,845 genes, as in the transductive benchmark; only the SL view was restricted to the inner-training labels.

Three initializations (456, 457 and 458) were fitted per arm and fold. The configuration used learning rate 0.001, hidden dimensions 512/256, neighborhood size 45, dropout 0.2, Alpha 2.0, Coe 2.0, Beta 4.0 and at most 300 epochs. The recorded training environment was TensorFlow 1.15.5, NumPy 1.21.6 and an A100-SXM4-80GB GPU. After each update, dropout was disabled for evaluation. Validation F1 was the maximum of $2PR/(P+R)$ over the precision–recall curve on validation positives and $V_{\mathrm{mix}}$, rather than F1 at a fixed probability threshold. Validation NDCG@10 ranked all 9,845 gene columns for each gene appearing in a validation-positive pair, masking training-positive pairs and using validation partners as relevant items. Thus outer-test genes could occur as unlabelled ranking candidates, but their labels were not used.

C1 selected the first epoch attaining the maximum $V_{\mathrm{mix}}$ F1 among trained epochs. The early-stopping counter reset whenever either this F1 or validation NDCG@10 strictly improved; training stopped after 20 epochs without either improvement once the zero-indexed epoch reached 100, ensuring at least 101 updates. NDCG affected stopping time but not C1's selection criterion. C2 fixed zero-indexed epoch 99 (100 updates), C3 used the final trained epoch, and C4/C5 selected the first maximum F1 on $V_{\mathrm{uni}}/V_{\mathrm{match}}$, respectively. All rules used the same trained trajectory.

Table S7 reports fold means after averaging the three initialization seeds. For Random training, C1 yielded T3 AUROC 0.5158 and a T0–T3 difference of 0.2504, satisfying the recorded thresholds of T3 $\leq 0.55$ and a decrease $\geq 0.15$. Across C1–C5, Random-trained T0 AUROC ranged from 0.7659 to 0.7729 and T3 from 0.5147 to 0.5210. The evaluation-population effect therefore persisted with outer-test-independent checkpoint selection. These seed-by-fold tables report benchmark discrimination. They do not contain SPIDR hit estimates for the validation-selected states.

**Table S7:** SLMGAE AUROC under outer-test-independent checkpoint selection. T1 uses the frozen full-graph matched set; T3 excludes held-out test positives from the degree calculation. These entries are means, not confidence intervals. Source: `E5_cv3_cells.csv`; seed/fold observations: `E5_cv3_per_fold.csv`.

| Training negatives | Checkpoint rule | T0 | T1 | T3 |
|---|---|---|---|---|
| Random | Validation F1 (C1) | 0.7662 | 0.4991 | 0.5158 |
| Random | Fixed epoch 99 (C2) | 0.7669 | 0.5043 | 0.5205 |
| Matched, training degree | Validation F1 (C1) | 0.7346 | 0.5104 | 0.5272 |
| Matched, training degree | Fixed epoch 99 (C2) | 0.7043 | 0.5295 | 0.5420 |

E3e numerical records are `E3E_summary_cells.csv` and `E3E_summary.json`.

### S10. Retrieval budget, regularization and SPIDR conditional analysis

E3c changed the evaluation geometry of the archived models to per-gene retrieval. SPIDR comprised 449 genes, with a median of 88 measured partners and budget $k = 10$. The ten negative redraws per design were summarized using a two-stage bootstrap over genes within fold and redraws. Random-trained LR retrieved 265.3 selected hits on average, compared with 232.8 for training-degree-matched LR and 182.7 expected under random ranking. Their respective gains were 1.840 pp [1.240, 2.469] and 1.116 pp [0.577, 1.727]; matched minus Random was −0.724 pp [−1.348, −0.093]. Full-graph matching gave a similar difference of −0.670 pp [−1.272, −0.016]. None of the 100 redraw pairings favored matched training at this budget. The HGB contrast was less conclusive: −0.448 pp [−0.953, 0.031]. SCHEMATIC, evaluated across 149 genes at $k = 3$, was also unresolved (LR difference −0.805 pp [−4.564, 3.065]).

These results establish a difference at a specified budget, not a budget-independent ranking of training designs. At SPIDR $k = 1$, matched LR gains were 3.35–3.48 pp versus 1.68 pp for Random. Moreover, each undirected pair appears in two per-gene lists; the E3c report notes that treating the gene lists as resampling units can produce intervals that are too narrow. The HAP1 degree-ranker comparator was calculated post hoc, and these shared models and evaluation pools do not constitute independent replications. Sources: `E3C_pergene_metrics.csv` and `E3C_primary_contrasts.csv`.

E3f aligned relative budgets because HAP1 top-10 represented approximately 0.5% of a candidate list, whereas SPIDR top-10 represented approximately 11%. At none of the three aligned fractional budgets did both screens have intervals excluding zero in opposite directions (Table S8). This met the recorded decision rule that budget could explain the original opposite-sign finding. It does not show that the screens are equivalent. A secondary descriptive sign rule relating the untrained degree ranker's gain to the training-design difference agreed in 11 of 12 HAP1/SPIDR screen–budget cells; those cells share data and are not independent external validations.

E3i varied LR regularization across $C = 10^{-4}, 10^{-3}, 10^{-2}, 0.1, 1, 10$, with ten negative redraws per design. The three primary values $(0.1, 1, 10)$ gave nearly identical HAP1 differences of approximately +0.74 pp and SPIDR differences of approximately −0.72 to −0.73 pp. At $C = 1$, the intervals were [0.32, 1.27] and [−1.33, −0.09], respectively. The strongest penalty retained the signs but

**Table S8:** LR retrieval-gain difference, training-degree matched minus Random, in pp with reported 95% intervals. Sources: `E3F_contrasts.csv` and `E3F_budget_grid.csv`.

| Budget | HAP1 | SPIDR |
|---|---|---|
| Top 10 | +0.74 [0.33, 1.27] | −0.72 [−1.35, −0.09] |
| Top 1% | +0.44 [0.19, 0.74] | +1.67 [−0.20, 3.59] |
| Top 5% | +0.13 [0.04, 0.23] | −0.53 [−1.44, 0.33] |
| Top 10% | +0.04 [−0.02, 0.10] | −0.68 [−1.30, −0.04] |

reduced the SPIDR contrast to −0.50 pp [−1.13, 0.10]. This supports stability within the examined regularization range, rather than invariance to arbitrary penalties. Sources: `E3I_C_grid.csv` and `E3I_contrasts.csv`.

### Conditional decomposition of SPIDR enrichment

E1 was explicitly a prespecified post hoc analysis: unconditional SPIDR enrichment had already been observed. For Random-trained SLMGAE forward scores, 21 hits occurred among 198 selections at the pooled top-1% budget. The whole-pool random expectation was 8.00. Redrawing selections within their joint endpoint-degree strata gave an expectation of 9.08, conditional enrichment of 2.31, and a one-sided permutation $p = 0.0003$ from 10,000 within-stratum permutations. Stratum composition accounted for 8.3% of the observed excess over the whole-pool expectation.

**Table S9:** SPIDR conditional selection expectations for the same 21 observed SLMGAE hits. Secondary permutation probabilities are nominal. Source: `E1_conditional_retrieval.csv`.

| Stratification | Conditional expectation | Observed/expected | One-sided $p$ |
|---|---|---|---|
| Joint endpoint-degree bins (primary) | 9.08 | 2.31 | 0.0003 |
| Degree-sum deciles | 9.31 | 2.26 | 0.0006 |
| Joint PubMed-count bins | 7.10 | 2.96 | 0.0001 |
| Annotation-richness deciles | 9.40 | 2.23 | 0.0006 |

In 200 constructed comparison-pool replicates, pairing the measured positives with degree-matched non-calls gave SLMGAE mean AUROC 0.560, with replicate 2.5th–97.5th percentiles [0.544, 0.576]. These are variation across constructed pools, not a confidence interval over biological experiments. Whole-pool SLMGAE AUROC was 0.594, close to degree alone (0.591), while the top-1% enrichment was 2.62-fold relative to uniform selection. The distinction between whole-ranking discrimination and head retrieval was therefore visible within the same measured pool.

Residual enrichment after these coarse covariates is not evidence of pair specificity. Eight of the 21 hits involved FZR1, and the strata did not preserve each individual gene's hit propensity. The analysis used one saved state per fold and retrospective full-graph degree information. Matched non-calls are a diagnostic comparison population; they are not verified biological negatives and do not define an experimentally achievable hit rate. Supporting metadata: `E1_summary.json`.

### S11. External isogenic screens and the Feng follow-up

#### Frozen test on six isogenic query backgrounds

E7b evaluated the Desjardins/Bowlan isogenic screens[11] after an eligibility audit and before model scoring. Six compatible loss-of-function backgrounds were retained: FBXW7, CDK12, ARID1A, KMT2D, STK11 and KEAP1. Candidate partners were mapped to the benchmark universe and restricted to the query's CV3 fold. The resulting pool contained 11,290 query–candidate entries, 63 positives under the authors' CCA top-50 definition, and 27 under the secondary top-20 definition. These are ranking-based hit definitions, not inferred statistical-significance thresholds.

The primary budget was the top 5% of each candidate list ($k = 94$ or 95), the smallest audited proportional budget with at least two expected random hits overall (3.16). Top-10 had only 0.33 expected hits and was secondary. The frozen sign rule predicted a Random-training advantage because the untrained degree ranker gained +0.32 pp. Primary LR matched-minus-Random gain was −0.20 pp [−0.78, 0.45]; HGB gave −0.30 pp [−0.83, 0.18]. Thus, the preregistered decision was weak consistency, not confirmation.

Intervals used 2,000 two-stage bootstrap resamples of the six queries and the negative redraws. Queries were resampled without fold stratification because four folds contained only one query. LR retrieved 7.7 hits with Random training and 6.6 with training-degree matching, versus 3.16 expected; respective one-sided probabilities from 2,000 within-query label permutations were 0.011 and 0.031. Excluding 41 known source pairs yielded a similar primary difference, −0.21 pp [−0.78, 0.41]. The secondary top-10% LR contrast excluded zero (−0.35 pp [−0.71, −0.03]), but does not replace the unresolved primary test. Six queries provide limited resolution, and variation across cell backgrounds is not separated from query identity here. Sources: `E7B_PREREG.md`, `E7B_geometry.csv`, `E7B_contrasts.csv`, and `E7B_hits_vs_random.csv`.

#### Feng screen: labels, direction, dose and conditional explanations

The Feng 2022 evaluation used BioGRID ORCS 2.0.18 BAGEL Bayes factors from 17 HEK293A genome-wide TKOv3 screens. Query screens 1845–1856 correspond, in order, to ARID1A, BAP1, CDH1, KEAP1, STK11, NF1, NF2, PBRM1, PTEN, RB1, TP53 and VHL; screen 1861 is the TP53BP1 query background. Screens 1857–1860 provide the four wild-type replicates. In each file, the `SCORE.1` column was used after deduplicating `IDENTIFIER_ID` (Entrez ID), retaining its first occurrence. The wild-type baseline was the per-gene mean BF across the four wild-type screens, skipping missing values. For each query, $\Delta$BF was query-background BF minus this wild-type mean, so larger values indicate greater dependency in the query background.

Genes represented in the query screen and wild-type baseline were ranked genome-wide by decreasing $\Delta$BF, with Entrez-ID strings in increasing order breaking ties (18,047 ranked genes per query). Ranking preceded mapping/filtering: numeric Entrez IDs were mapped to the 9,845-gene benchmark universe, the query itself and unmapped genes were removed, and partners were restricted to the query's CV3 fold. Duplicate mapped partners retained their earliest rank. Primary and secondary positives had genome-wide ranks $\leq 50$ and $\leq 20$, respectively. The resulting 13 query lists contained 24,909 candidate entries, 115 top-50 positives and 50 top-20 positives, with 1,909–1,934 candidates per list. These are operational differential-dependency rank labels, rather than significance thresholds or independently validated SL interactions. The primary budget was $k = \lceil 0.05n \rceil$; secondary budgets were $k = 10$ and $\lceil 0.10n \rceil$.

The untrained full-SynLethDB-degree ranker gained +1.149 pp at the primary budget, predicting a Random-training advantage under the prespecified sign rule. Ten negative redraws per design yielded the results in Table S10. E8a contrasts used 2,000 two-stage bootstrap replicates (seed 20261024): the 13 queries were resampled as one group without fold stratification, because one fold contained only a single query, and the ten redraw indices were sampled independently for each training arm. Intervals are the 2.5th and 97.5th percentiles. LR matched-minus-Random was −0.656 pp [−1.146,−0.174], consistent with the predicted direction. Removing 296 known SynLethDB pairs preserved the difference (−0.62 pp [−1.13,−0.16]); omitting the ARID1A knockdown query gave −0.57 pp [−1.10,−0.09]. The pooled Feng/E7b estimate across 19 queries was −0.51 pp [−0.93,−0.12]; this overlaps both constituent evaluations and is not another independent replication.

**Table S10:** Feng top-5% retrieval with the top-50 differential-BF label. Hits are means over ten negative redraws, summed over query lists; the exact uniform-ranking expectation is 5.77. Differences are in pp with 95% two-stage percentile intervals. Sources: `E8A_contrasts.csv`, `E8A_results_by_model.csv`, and `E8A_coupling.csv`.

| Learner | Random hits | Matched hits | Gain difference [95% interval] | Coupling, R → M |
|---|---|---|---|---|
| LR | 16.7 | 8.5 | −0.66 [−1.15, −0.17] | 0.786 → 0.184 |
| HGB | 16.4 | 7.0 | −0.75 [−1.18, −0.37] | 0.757 → 0.297 |

Random-trained LR and HGB each exceeded the within-query label-permutation null ($p = 1/2001 \simeq 0.00050$). Training-degree-matched LR and HGB did not show resolved gains against this uniform-ranking null ($p = 0.1284$ and 0.2309). The sign rule agreed in ten of twelve Feng label–budget–learner cells; both exceptions used $k = 10$. These cells share queries and models.

E8b varied the matched-negative fraction over $0, 0.25, 0.5, 0.75, 1$, with five redraws per dose. LR coupling decreased from 0.787 to 0.186; its mean Feng gains were 0.848, 0.720, 0.656, 0.384 and 0.193 pp, with 16.4, 14.8, 14.0, 10.6 and 8.2 hits. Spearman dose associations were −0.981 for coupling and −0.947 for gain; HGB gave −0.973 and −0.887. Each one-sided probability was 1/10001, using 10,000 permutations of dose labels across the 25 dose–redraw observations (seed 20261024). On Random negatives, removing 28 degree features reduced LR gain by 0.159 pp; retaining only the 17 jaccard/edge/neighbor-jaccard features reduced it by 0.512 pp. Matched-training changes were +0.032 and +0.144 pp. These interventions share the measured pool and are not independent biological replications. Sources: `E8B_dose_points.csv`, `E8B_trend_tests.csv`, and `E8B_ablation.csv`.

E8c exchanged 28 endpoint coefficients and 31 pair-feature coefficients on the original feature scale, pairing Random and matched models by redraw index as in S8. Queries were resampled without fold stratification, but the ten redraw-pair indices were sampled jointly across endpoint, pair, total and interaction effects (2,000 replicates, seed 20261024). The total LR contrast was −0.656 pp [−1.168,−0.183]; the endpoint effect was −0.508 pp [−0.897,−0.156], and the pair effect was −0.147 pp [−0.358,0.056]. The endpoint effect had the same sign as the total, excluded zero and represented 78% of its point estimate, satisfying the recorded endpoint-effect criterion. This coefficient-block decomposition is not a biological mediation fraction. E8a and E8c have the same total point estimate but different intervals because independent-arm redraw resampling in E8a differs from joint model-pair resampling in E8c. Source: `E8C_factorial.csv`.

E8d used within-fold checkerboard swaps preserving each query's and partner gene's hit counts, with fitted scores fixed. For a fold containing $m$ positives, burn-in comprised $20m$ attempted swaps and consecutive saved samples were separated by $5m$ attempts, yielding 2,000 null samples. The observed LR matched-minus-Random hit difference was −8.2, compared with a null mean of −8.886 (SD 1.748; one-sided $p = 0.6637$). Random-trained LR had 16.7 hits versus conditional expectation

18.07; matched LR had 8.5 versus 9.18. The ratio of null mean difference to observed difference was 108%, a point ratio without an interval. These results are compatible with gene-level hit propensities accounting for the design difference; they do not prove the absence of pair-specific information or identify a causal explained fraction. One fold had a single query and could not be randomized. In the swappable folds, acceptance rates were 0.491–0.716 and mean positive-retention fractions 0.247–0.509; these summaries do not establish complete chain mixing. Conditional expectations are distinct from the uniform-ranking expectation of 5.77 hits. Sources: `E8D_null_tests.csv` and `E8D_chain_diagnostics.csv`.

### S12. MLP replication and limits of endpoint ablation

**Architecture, fitting and endpoint-input shuffle.** The feature MLP used two hidden layers of 64 and 32 ReLU units and a logistic binary output, L2 penalty $\alpha = 10^{-4}$, at most 200 training epochs, and `random_state`$= 20261025 + r$ for zero-indexed redraw $r$. Missing features were filled with medians from the arm's source-positive/negative training set, then transformed by a StandardScaler fitted to that same set; evaluation pools used those fitted medians and scaling parameters. Scores were positive-class probabilities. Early stopping was enabled with a 10% training-data validation fraction and `n_iter_no_change=10`. This MLP validation split is an internal pair-level split, unlike E5's gene-isolated validation design.

The remaining `MLPClassifier` arguments used library defaults. The reference specification checked against scikit-learn 1.9.1 uses Adam with initial learning rate 0.001, $\beta_1 = 0.9$, $\beta_2 = 0.999$, $\epsilon = 10^{-8}$, automatic batch size $\min(200, n)$, sample shuffling, and tolerance $10^{-4}$. Its early-stopping validation split is label-stratified, assessed by accuracy, and restores the best validation weights. The archived run did not record the scikit-learn version, so 1.9.1 identifies the default-settings reference, not a verified historical runtime. E9 comprised 125 all-feature fits (five doses, five redraws, five folds) and 50 A1 fits (two endpoint doses, five redraws, five folds), totaling 175 fits.

For the frozen-model shuffle, each pair's entire 28-dimensional endpoint vector was permuted as one block; the 31 pair-feature columns stayed fixed. The block comprises minimum/maximum degree features for both endpoints, so this is not simply a shuffle of the partner gene. Permutations were applied across all test pairs in a fold-specific screen pool, including across HAP1 queries within that fold, rather than within each query. For each fitted model, a generator with seed $20261024 + 100k + 7$ (zero-indexed fold $k$) generated one row permutation for each pool in the order HAP1, SPIDR, SCHEMATIC, Feng. The generator was reset identically for every fit, giving the same permutation within a fold/pool across redraws and training arms. All-feature models at doses 0 and 1 were rescored without refitting using their original medians, scalers and weights.

An MLP trained on the same 59 KG features was evaluated over two negative designs, ten redraws and five folds (100 fits; E5-MLP). HAP1 top-10 gain increased from 0.188 to 0.529 pp, a difference of 0.341 pp with reported interval [0.000, 0.762]. The unrounded lower bound was $4.28 \times 10^{-17}$, numerically zero, so the primary HAP1 result provides partial rather than resolved support. HAP1 top-5 and top-5% differences were 0.476 pp [0.061, 1.061] and 0.081 pp [0.012, 0.165]. SPIDR top-10 favored Random training, with difference −0.650 pp [−1.145, −0.127]. HAP1 score–degree coupling decreased from 0.534 to 0.128, difference −0.406 [−0.441, −0.372]. Sources: `E5_MLP_results.csv` and `E5_MLP_summary.json`.

E9a then evaluated five matched-negative fractions with five redraws per dose. Table S11 distinguishes the primary mean coupling endpoint from the retrieval screens: the stored `coup_mean3` is the mean of HAP1, SPIDR and SCHEMATIC coupling, not a mean including Feng. The coupling trend was negative ($\rho = -0.977$, one-sided $p \simeq 0.0001$). HAP1 gain increased overall ($\rho = 0.567$,

$p = 0.0020$), SPIDR gain decreased ($\rho = -0.530$, $p = 0.0053$), and the secondary Feng trend decreased ($\rho = -0.696$, $p \simeq 0.0001$), using 10,000 dose-label permutations over the 25 dose–redraw observations. HAP1's mean curve was not monotonic and its endpoint change was approximately 0.30 pp, smaller than the corresponding LR change of approximately 0.74 pp. Feng also rose between doses 0.75 and 1; the rank-trend test is not a claim of strict monotonicity.

**Table S11:** MLP dose-response means over five negative redraws per dose. Retrieval gains are in pp. Mean coupling averages HAP1, SPIDR and SCHEMATIC; Feng coupling is separate. Sources: `E9_metrics.csv` and `E9A_trend_tests.csv`.

| Matched fraction | 0 | 0.25 | 0.50 | 0.75 | 1 |
|---|---|---|---|---|---|
| Mean coupling | 0.548 | 0.498 | 0.414 | 0.307 | 0.086 |
| HAP1 gain | 0.18 | 0.23 | 0.51 | 0.41 | 0.49 |
| SPIDR gain | 1.55 | 1.50 | 1.30 | 1.14 | 1.01 |
| Feng gain | 0.67 | 0.43 | 0.18 | 0.05 | 0.16 |
| Feng coupling | 0.605 | 0.578 | 0.511 | 0.394 | 0.180 |

E9b assessed two point-estimate criteria across HAP1, SPIDR and Feng: retrained A1 ablations had to follow the predicted directions and be smaller in magnitude under matched than Random training, and frozen-model shuffling had to remove at least half the Random–matched coupling gap in each screen. Shuffling Random-trained MLP endpoint inputs reduced HAP1/SPIDR/Feng coupling from 0.543/0.497/0.605 to 0.239/0.177/0.334, removing 74%/63%/64% of the respective gaps. Retrieval gain changed by $-0.05/-0.89/-0.50$ pp. The shuffle criterion therefore held, supporting a contribution of endpoint inputs to the fitted score dependence.

The retraining criterion did not hold. Removing degree features under Random training changed HAP1/SPIDR/Feng gain by $+0.134/-0.290/+0.016$ pp; the small Feng increase was opposite to the predicted decrease. Under matched training, the respective changes were $+0.329/+0.053/+0.143$ pp, so the HAP1 and Feng changes were larger than under Random training. Lower marginal score–degree coupling therefore did not imply that a matched MLP ceased using explicit degree features. A reversed use of degree by the matched model remains an untested explanation. E9b's partial-support conclusion is based on the two point-estimate criteria; its ablation/shuffle assessment did not specify confidence intervals. Sources: `E9B_ablation.csv` and `E9_metrics.csv`.

### S13. HuRI cross-task test and refutation of the directional rule

**Universe, candidate lists and reference labels.** The PPI follow-up used the 9,845-gene benchmark universe, mapped one-to-one by Ensembl gene ID after removal of version suffixes, and the same five-fold node split (`KFold(5, shuffle=True, random_state=123)`). Lit-BM-17 supplied source positives: 7,665 non-self, unique unordered pairs remained within this universe. Training positives had both endpoints outside the test fold, yielding 4,955, 5,013, 4,753, 4,766 and 5,000 pairs across folds.

The operational candidate universe was all unordered within-fold pairs among genes belonging to HuRI Space III (`in_space_3=1` in Supplementary Table 1), not a pair-by-pair assay-completion mask. Of 17,408 Space III genes, 9,243 mapped to the benchmark universe. Their fold counts were 1,837, 1,855, 1,845, 1,857 and 1,849, defining 8,538,813 candidate pairs in total. HuRI[12] calls labelled 3,061 of these pairs positive; the secondary HI-union definition labelled 3,839. Noncalled pairs served as negative references within this operational universe. Gene-level Space III membership does not establish that every pair was successfully measured or that an uncalled pair lacks an interaction; assay coverage and limited Y2H sensitivity can therefore affect this comparison.

Primary retrieval evaluated the 2,111 genes with at least one within-fold HuRI-positive partner (391, 444, 462, 428 and 386 by fold). Each query ranked every other Space III gene in its fold, with median candidate-list size 1,848. The implementation stored lists for 2,424 genes with a HuRI or HI-union positive partner and selected the relevant query subset for each label definition. Selected-hit totals count incidences across lists, not necessarily unique interactions. The main pool retained source-positive pairs within the test fold; a sensitivity analysis excluded every Lit-BM pair, and the HuRI-novel analysis removed HuRI positives flagged as appearing in the earlier Rual, Venkatesan, Yu, Rolland or Yang screens in Supplementary Table 11.

**Features, learners and negative sampling.** The 54 symmetric KG features excluded SL, non-SL, SR and the direct interaction relation `INTERACTS_GiG`. Eleven annotation relations (GO BP/MF/CC, pathways, expression, disease association, disease-associated up/downregulation, compound binding, and compound-associated up/downregulation) each contributed log-transformed shared-neighbor count, Jaccard overlap and log-transformed minimum/maximum endpoint degree, for 44 columns. The regulation and covariation gene–gene relations each contributed an edge indicator, log-transformed common-neighbor count, neighbor Jaccard overlap and the two degree columns, for ten more. The endpoint block contained 26 columns and the pair block 28; DEGIDX was the mean of the endpoint columns. A1 retained the 28 pair columns, A2 the 15 jaccard/edge/neighbor-jaccard columns, and the no-GO sensitivity removed 12 GO columns, leaving 42 features.

LR used training-median imputation and StandardScaler, $C = 1$, `lbfgs` and at most 5,000 iterations. HGB used 200 iterations, learning rate 0.1, at most 31 leaves, minimum leaf size 20, no L2 penalty or early stopping, and seed 20261024. Both ranked their decision-function scores. The MLP used the configuration in S12 and positive-class probabilities.

Training negatives were sampled 1:1 to positives from all 7,876 outer-training genes, not only genes observed in Lit-BM or Space III. Self-pairs, duplicates and all 7,665 known Lit-BM pairs were excluded. Random sampling drew uniform endpoints with rejection. Both matched designs used the larger/smaller endpoint bins $\lfloor \log_2(d+1) \rfloor$ and the cell-matching/fallback algorithm described in S9; full-source matching used all Lit-BM degrees, whereas training matching used only that fold's training positives. The ten negative-redraw seeds were 1001–1010, 2001–2010 and 3001–3010 for Random, full-source matched and training-degree matched, with one generator consumed sequentially across folds for each redraw. At intermediate matched doses, the mixture seeds were $4000 + 100j + 10r + k$ for zero-indexed dose-grid, redraw and fold indices; the endpoint doses reused the first five base redraws.

**Primary contrast and uncertainty.** LR was primary, with HGB and MLP additional learners. E10a used 2,000 two-stage bootstrap replicates (seed 20261024): query units were resampled with replacement within their folds, while each arm's ten redraw indices were resampled independently. Each replicate first averaged across its sampled redraws and then its sampled queries; 95% intervals are the 2.5th and 97.5th percentiles. Query lists can share an underlying pair, so this interval describes the specified query-resampling procedure rather than independent-edge or independent-biological-experiment uncertainty.

The frozen predictor ranked candidates by partner degree in the complete Lit-BM graph. Its HuRI gain was +0.260 pp, so the rule predicted that Random training would outperform matched training. The observed differences instead favored matched training for all three learners, with intervals excluding zero (Table S12). The prespecified primary decision was therefore refutation of the sign rule, not confirmation, even though matched training improved retrieval.

**Table S12:** HuRI top-10 results; matched denotes training-graph matching. Gain differences are in pp. Uniform random expectation was 33.13 selected hits. Sources: `E10A_contrasts.csv`, `E10A_results_by_model.csv`, and `E10A_per_fold.csv`.

| Learner | Random hits | Matched hits | Gain difference [95% interval] |
|---|---|---|---|
| LR | 226.7 | 288.2 | +0.291 [0.191, 0.392] |
| HGB | 221.7 | 279.7 | +0.275 [0.193, 0.361] |
| MLP | 223.3 | 276.6 | +0.252 [0.158, 0.351] |

For LR, absolute gain was 0.917 pp [0.774, 1.058] under Random training and 1.208 pp [1.043, 1.375] under training-graph matching. All 15 learner–fold differences were positive. Sensitivity analyses crossed two label definitions (HuRI and HI-union), three budgets (top-10, top-5%, top-10%) and three pools (unrestricted, source-pair excluded, HuRI-novel), with one additional minimum-positive-count analysis. Across 19 cells per learner and two matched contrasts, all 114 differences were positive with reported intervals excluding zero. These are correlated sensitivity analyses, not 114 independent confirmations. LR remained positive after source-pair exclusion (+0.24 pp [0.15, 0.33]), on HuRI-novel pairs (+0.20 pp [0.12, 0.30]) and using HI-union labels (+0.24 pp [0.14, 0.33]).

E10b used five matched-negative fractions and five redraws per fraction. LR coupling changed from 0.772 to −0.546, while HuRI gain rose from 0.92 to 1.21 pp; intermediate gains were 1.13, 1.17 and 1.22 pp. HGB and MLP gains changed from 0.90 to 1.17 and from 0.91 to 1.16 pp, respectively. Coupling decreased with dose for all learners ($\rho = -0.981, -0.981, -0.973$; each one-sided permutation $p \simeq 0.0001$). However, retrieval moved opposite to its predicted decrease: $\rho = +0.899, +0.791, +0.638$, with one-sided probabilities for the prespecified negative trend of 1.000, 1.000 and 0.9997. This directional test failed.

The ablation result also opposed the original prediction. In Random-trained LR, removing 26 endpoint features increased HuRI gain by 0.03 pp, and retaining only 15 jaccard/edge/neighbor-jaccard features increased it by 0.24 pp; matched-training changes were −0.03 and −0.01 pp. The descriptive pattern agrees with the observed matched advantage but does not convert the failed prediction into confirmation. Sources: `E10B_dose_points.csv`, `E10B_trend_tests.csv`, and `E10B_ablation.csv`.

The cross-task evidence therefore separates two claims. Changing negative sampling altered degree coupling in PPI as well as SL, but the sign of the untrained source-degree ranker's retrieval gain did not correctly predict the PPI training-design comparison. Any proposed directional rule must retain this counterexample and be restricted to its tested SL settings and budgets.

**PPI weight swaps and conditional null.** E10c exchanged the LR endpoint-coefficient block without refitting, after conversion to the original feature scale as in S8. Random and matched models were paired by redraw index. Its 2,000 bootstrap replicates (seed 20261024) resampled the 2,111 queries within folds and the ten model-pair indices jointly across endpoint, pair, total and interaction effects, with 95% percentile limits. This joint effect resampling differs from E10a's independent-arm redraw resampling and explains the small difference between their total-contrast intervals. The primary endpoint effect was 0.297 pp [0.208,0.384], compared with a pair-block effect of −0.006 pp [−0.050,0.040] and total contrast 0.291 pp [0.192,0.393]. The endpoint-to-total point ratio was 102%, reflecting a small offsetting pair-block effect. Norm-preserving swaps gave an endpoint effect of 0.204 pp [0.127,0.283]; full-source matching gave 0.291 pp [0.203,0.378]. They support a functional coefficient-block comparison within this representation, rather than biological mediation. Source: `E10C_factorial.csv`.

E10e retained fitted rankings and rearranged HuRI positive edges by within-fold degree-preserving double-edge swaps. Across 2,000 saved null draws, observed training-degree-matched-minus-Random hit-incidence differences were 61.5 (LR), 58.0 (HGB) and 53.3 (MLP), compared with null means 5.30, 5.23 and 5.63 (SD 5.86, 4.13 and 4.67). Each one-sided exceedance probability was 1/2001. For LR, the absolute hit totals were 226.7 and 288.2, versus conditional expectations 32.05 and 37.35. Acceptance rates were 0.884–0.936 and mean edge-retention fractions 0.031–0.048. Both the absolute rankings and the design differences therefore exceed this fixed-degree null; the null expectations are not identified causal shares. Sources: `E10E_null_tests.csv` and `E10E_chain_diagnostics.csv`.

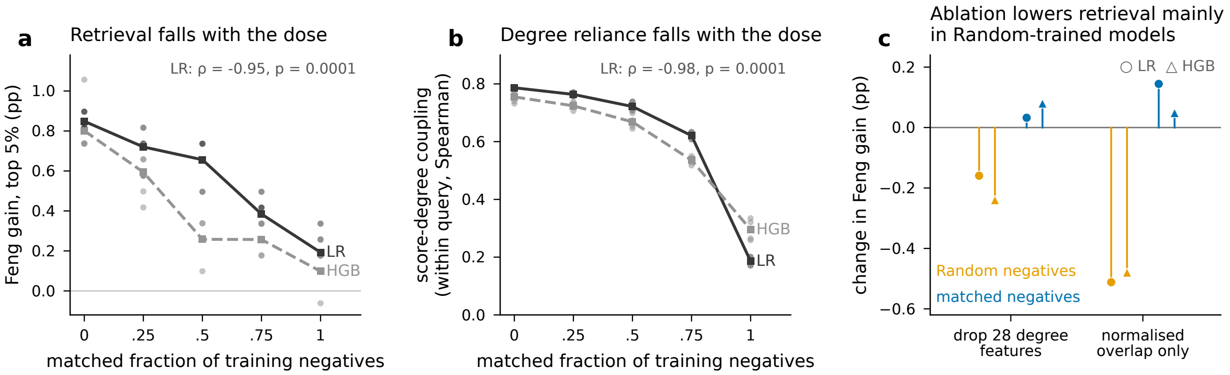


**Figure S3: Sampling-dose response in the Feng panel.** Five negative redraws per dose; LR and HGB show decreasing coupling and overall retrieval trends as the matched fraction rises. Dots are redraws and squares their means, not confidence intervals. LR's retrieval trend is $\rho = -0.947$ (one-sided dose-label permutation $p \simeq 0.0001$). Panel c shows feature-ablation changes relative to all-feature models. Trend probabilities use 10,000 dose-label permutations within the fixed-pool analysis.

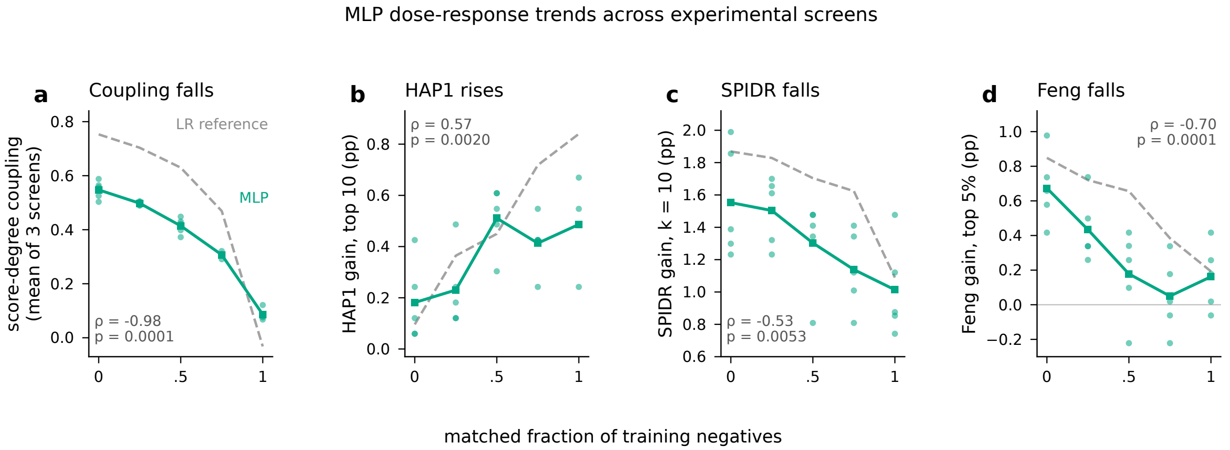


**Figure S4: MLP dose-response extension.** Dots show five negative redraws per dose and squares their means, not confidence intervals. Grey dashed LR series in panels a–c come from E3d and the Feng series in panel d from E8. Mean coupling averages HAP1, SPIDR and SCHEMATIC. Feng retrieval has an overall negative rank trend ($\rho = -0.696$, one-sided dose-label permutation $p \simeq 0.0001$), although its mean rises from 0.050 to 0.162 pp between doses 0.75 and 1; HAP1 is also not monotone at every dose. The companion retraining ablation did not satisfy the predicted direction/asymmetry criterion, while endpoint-input shuffling removed more than half the coupling gap in each screen.

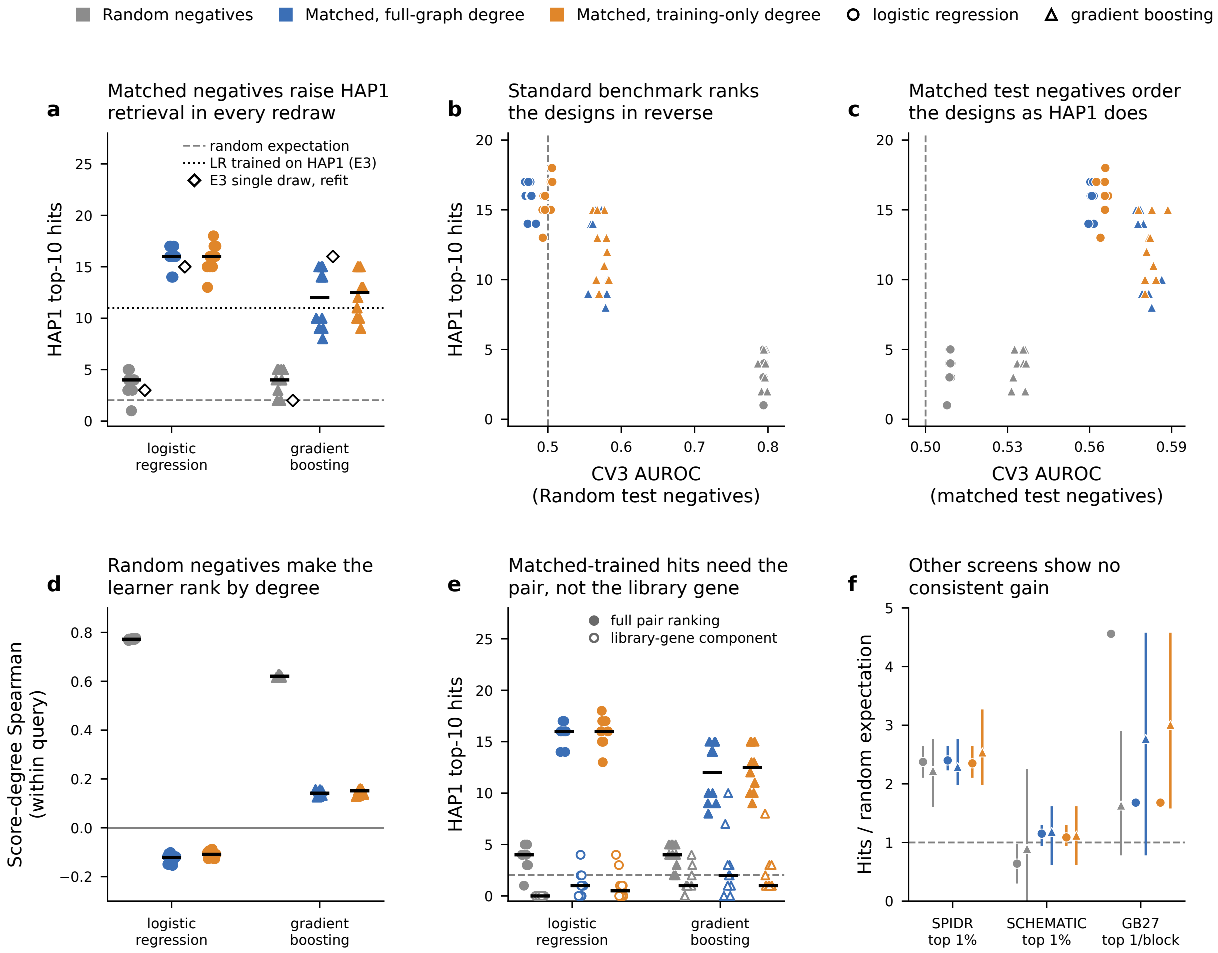


**Figure S5: Changing training negatives changes HAP1 retrieval.** Ten redraws per design retain the same source positives and representation. (a) HAP1 top-10 hits over 164 queries; black ticks denote medians, diamonds the preceding single-draw fits, and the dotted reference the separately HAP1-trained LR. (b,c) CV3 AUROC under Random or matched test negatives against HAP1 hit counts. (d) Within-query score association with the knowledge-graph endpoint-degree summary. (e) Full-model retrieval and a library-gene-only score component. (f) Other experimental pools, shown as hits relative to each pool's random expectation; markers are means and bars are redraw ranges. Redraws reuse the same experimental labels. GB27 contains five effective pair selections repeated across cell lines.

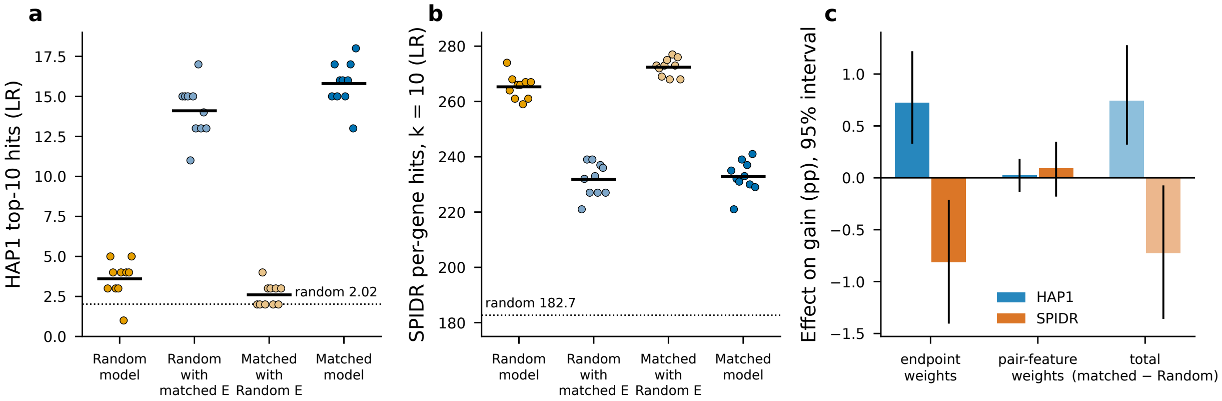


**Figure S6: Endpoint-coefficient swaps transfer the retrieval contrast.** (a,b) LR retrieval before and after exchanging only the 28 endpoint coefficients; the remaining pair-feature coefficients retain their indicated training origin. Points represent ten paired negative redraws; horizontal bars show means. The dotted lines are task-specific random-ranking expectations. HAP1 uses 164 query top-10 rankings; SPIDR uses 449 gene top-10 rankings, so counts in (b) are retrieval incidences rather than unique pairs. (c) Symmetric factorial endpoint and pair-block effects and the total matched-minus-Random contrast, in precision-gain percentage points with two-stage 95% bootstrap intervals. The blocks are correlated sources of model information; effect ratios are not causal mediation fractions.

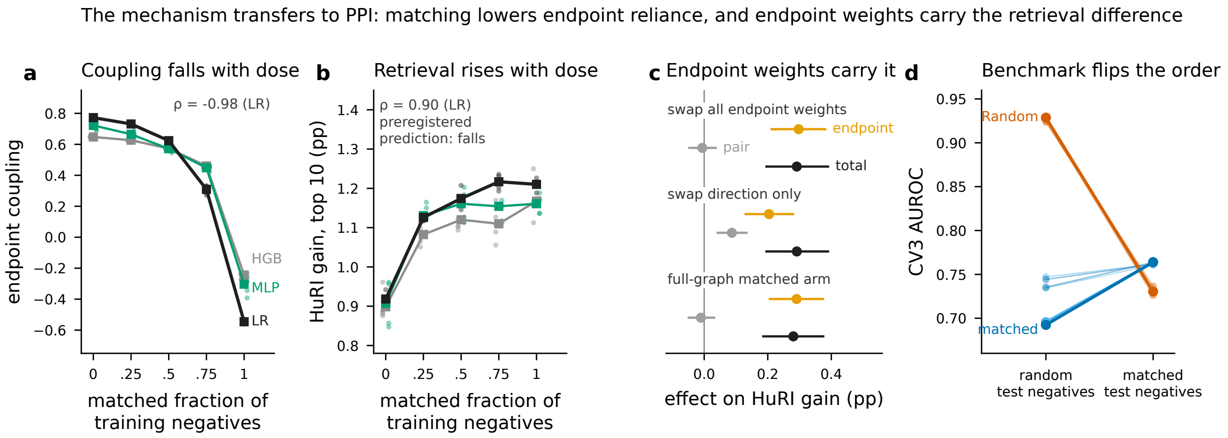


**Figure S7: Endpoint reliance and retrieval in the PPI extension.** (a,b) Increasing the matched fraction reduces score–endpoint coupling while improving HuRI retrieval, opposite to the prespecified retrieval-direction rule. Points show redraw values and lines their means for LR, HGB and MLP. (c) LR coefficient-swap effects in precision-gain percentage points, with two-stage 95% bootstrap intervals: exchanging all endpoint coefficients, preserving their recipient norm, or using full-source matching. (d) LR CV3 discrimination under Random and matched tests. The transferable effect concerns this model's use of endpoint attributes; the failed direction prediction is retained. HuRI retrieval counts are query-ranking incidences, not independent pair discoveries.

### S14. Degree-preserving rewiring of source-positive pairs

**Intervention and fixed quantities.** This control changes the source-positive partner assignments, rather than the fitted coefficients or experimental labels. It uses the official five-fold CV3 split and the first five archived negative redraws in their original order, with no outcome-based selection. For each fold and redraw, one chain starts from the original source-positive graph and produces a graph shared by the Random and matched-negative training arms. Both endpoints of every source edge remain within that fold's training genes. The forbidden-edge set is the union of the two fixed negative lists for that fold and redraw. The original positives may be retained or reappear; they are not permanently forbidden.

The undirected simple-graph sampler proposes symmetric double-edge switches for a fixed $50m$ attempted steps, where $m$ is the source-positive edge count. Rejected proposals count toward this total. Self-loops, duplicate edges and forbidden negative edges are rejected. The procedure preserves the number of edges and each gene's positive degree exactly. Original-edge retention and acceptance are recorded at $m$, $5m$, $10m$, $25m$ and $50m$ attempts. The predefined diagnostic requires replacement of at least half the original edges and a change in retention below 0.02 between $25m$ and $50m$. These checks assess movement and late-chain stability; they do not prove uniform sampling, irreducibility or complete mixing under the forbidden-edge constraints. Degree-preserving training controls have precedents in interaction prediction [14]; the present comparison concerns measured external retrieval with fixed negative lists.

The 59-feature representation and learner settings remain fixed. LR uses training-median imputation, StandardScaler, $C = 1$, lbfgs and at most 5,000 iterations. HGB uses 200 iterations, learning rate 0.1, at most 31 leaves, at least 20 observations per leaf, zero L2 penalty, no early stopping and random state 20261024. There are 25 rewired graphs and 100 new fits (five folds, five redraws, two negative lists and two learners). LR's original-positive references were refitted in the same runtime before comparison, after a reference check detected differences from the archived predictions; 50 reference fits use unchanged data and hyperparameters. HGB references reuse archived predictions after exact checks of scores and top-10 inclusion weights for both arms in redraw 1, fold 1. The reference-handling amendment preceded generation of the rewired graphs and outcomes. The current runtime used NumPy 2.4.6 and scikit-learn 1.9.1.

**Evaluation and uncertainty.** HAP1 uses its frozen 309,372 measurement rows, 164 query lists and primary label A; SPIDR uses 19,664 primary unordered pairs and 449 query genes with at least 20 candidate partners. Both tasks here use top 10 per query, with the archived fractional inclusion rule for boundary ties. SPIDR totals therefore count query-ranking incidences and differ from the earlier fold-global top-1% task. Gain is query precision minus the query's candidate prevalence, averaged across queries and then across redraws. Source-pair effects are $G_{\text{original}} - G_{\text{rewired}}$; positive values indicate reduced retrieval after rewiring.

Paired two-stage bootstrap samples resample the five complete redraws and queries within their official folds. The same resampling indices are used for both source graphs, negative lists and learners (10,000 samples). We report descriptive 95% percentile intervals and, for the eight source-pair contrasts, per-cell 99.375% intervals implementing a Bonferroni family-95% adjustment. This family does not cover every historical analysis in the project. Repeated draws and folds are computational comparisons, not independent biological repetitions. Shared partners and repeated pair incidences, particularly in SPIDR, limit the independence assumptions of query resampling. Per-fold and leave-one-fold-out summaries are descriptive sensitivity checks.

In the fixed-M comparison, HAP1 LR mean hit incidences decrease from 15.8 to 4.4 and HGB

**Table S13:** Source-positive rewiring: all eight comparisons. Gains and original-minus-rewired differences are percentage points (pp). Fixed M denotes the original training-degree-matched negative list, retained after positive rewiring. The final column gives the Bonferroni-adjusted bootstrap interval for the difference. Source: `rewiring/analysis/summary_8cells.csv`.

| Pool | Model | Negatives | Original | Rewired | Difference | Adjusted interval |
|---|---|---|---|---|---|---|
| HAP1 | LR | Random | 0.096 | -0.075 | 0.171 | [-0.110, 0.573] |
| HAP1 | LR | Fixed M | 0.840 | 0.145 | 0.695 | [0.113, 1.488] |
| HAP1 | HGB | Random | 0.133 | -0.014 | 0.146 | [-0.192, 0.524] |
| HAP1 | HGB | Fixed M | 0.584 | 0.035 | 0.549 | [0.040, 1.265] |
| SPIDR | LR | Random | 1.873 | 1.766 | 0.107 | [-0.503, 0.698] |
| SPIDR | LR | Fixed M | 1.089 | 0.711 | 0.379 | [-0.298, 1.090] |
| SPIDR | HGB | Random | 1.597 | 1.138 | 0.459 | [-0.311, 1.243] |
| SPIDR | HGB | Fixed M | 1.103 | 0.461 | 0.641 | [-0.091, 1.359] |

from 11.6 to 2.6, against 2.023 expected under uniform ranking. These HGB counts concern the first five redraws; the ten-redraw comparison in Section S8 gives 12.3 matched-training hits and should not be substituted here. The two source-pair effects remain positive in all five redraw summaries, all five folds and all five leave-one-fold-out summaries. The third fold contributes substantially, but removing it does not reverse either effect. SPIDR Random-trained LR retains a positive rewired gain of 1.766 pp (95% interval 1.194–2.359), versus 1.873 pp with original positives. The unresolved difference does not establish equivalence. Different significance outcomes between pools do not constitute a formal test of cross-pool heterogeneity.

The supporting interaction is $(G_M - G_R)_{\text{original}} - (G_M - G_R)_{\text{rewired}}$. It is 0.524 pp [0.159, 0.976] for HAP1 LR and 0.402 [0.061, 0.829] for HAP1 HGB. SPIDR estimates are 0.272 [$-0.334$, 0.886] and 0.183 [$-0.486$, 0.829], respectively. These are descriptive 95% intervals, not additional members of the eight-contrast corrected claim.

**Structural diagnostics and scope.** All 25 graphs preserve every node's positive degree and pass the partition, duplicate, self-loop and forbidden-edge checks. Original-edge retention is 7.39–11.09%, acceptance is 56.70–64.01%, and the maximum late retention change is 0.00552. All 100 new fits complete with finite, nonconstant predictions. Rewired training AUROC ranges are 0.797–0.818 (LR Random), 0.532–0.545 (LR fixed M), 0.925–0.932 (HGB Random) and 0.800–0.833 (HGB fixed M).

Marginal degree preservation does not preserve joint endpoint-bin frequencies, assortativity, endpoint minima/maxima or pair-feature distributions. The total-variation distance between original and rewired joint source-degree-bin distributions is 0.286–0.324. Consequently, the fixed-M lists are no longer guaranteed to match the rewired positives' joint bins. The fixed-negative comparison measures the joint effect of changing source partner assignments and their degree-bin alignment with the negative list. It cannot distinguish these two changes. Rewired edges are not experimentally established negatives, some original edges remain, and auxiliary graph features retain biological information. Historical target labels and effects were already known; freezing this analysis and reading labels after predictions does not make it a blinded biological experiment. Full feature-shift, per-redraw and per-fold tables are retained with the source results.

### S15. Predicting retrained model behavior from negative distributions

**Density-ratio predictor.** For each source task, fold and negative redraw, a discriminator labels Random negatives as 0 and matched negatives as 1. Class sizes are equal; source positives, fitted learner outputs and target labels are not discriminator inputs. Training-median imputation and StandardScaler are fitted to the combined negative data. A logistic discriminator selects $C \in \{0.001, 0.01, 0.1, 1, 10\}$ by minimum out-of-fold log loss under stratified five-fold splitting (shuffle, seed 20261027), choosing the smaller $C$ on a tie. It is then refitted to all negative samples using lbfgs and at most 5,000 iterations. Its logit estimates $\log \widehat{r}(x)$, where $r(x) = M(x)/U(x)$ in the observed feature space. Coefficients are converted back to the original feature scale. The log-ratio is clipped at the 0.5th and 99.5th percentiles of that discriminator's combined source-negative logits.

The prediction is $\widehat{s}_M = s_R - \mathrm{clip}(\log \widehat{r})$. The fitted matched learner supplies the target behavior to predict, not an input to this prediction. LR and HGB use decision-function scores; the MLP comparison transforms clipped probabilities to logits (clipping limits $10^{-6}$ and $1 - 10^{-6}$). SL models retain 59 features; PPI models retain 54 after removal of the direct interaction relation. The primary comparison uses ten Random/matched redraw pairs per fold, with M_train matching based on the source training graph. M_full matching, untruncated ratios, endpoint-only and pair-only discriminators, a one-dimensional degree discriminator and a Random-trained endpoint-ablated model are sensitivity or comparator analyses. MLP comparisons use five archived redraws. The secondary HGB discriminator uses the same 200-iteration HGB configuration, fitted once to all source negatives without internal hyperparameter selection; probabilities are clipped to $[10^{-6}, 1 - 10^{-6}]$ before taking logits and applying its own source-derived percentile limits.

The source-only feasibility checks gave mean discriminator cross-validation AUROCs of 0.805 for SL and 0.885 for PPI. The proportion of target candidates inside the source-derived clipping interval was 0.996 (HAP1), 0.998 (SPIDR), 0.993 (Feng) and 0.992 (HuRI's full candidate universe). Passing these checks establishes usable feature-space separation and range coverage, not transportability of a biological label distribution. The matched models and screen results had been seen before E11; this is a prespecified retrospective behavior-prediction analysis.

**List-overlap endpoint (P1).** Let $\pi_A(i)$ be candidate $i$'s inclusion weight in a top-$k$ list from score $A$, including fractional boundary ties. Overlap is $O(A, B) = \sum_i \pi_A(i)\pi_B(i)/k$. Within each redraw, overlaps are averaged over lists before computing

$$C = \frac{\overline{O}(\widehat{s}_M, s_M) - \overline{O}(s_R, s_M)}{1 - \overline{O}(s_R, s_M)}.$$

The primary statistic averages this closure over redraws. A mean denominator below 0.05 makes a pool non-estimable. The primary criterion requires mean closure across the four estimable pools of at least 0.5 and a positive lower 95% interval bound in every estimable pool. A per-list-ratio calculation is secondary; it is not substituted for the primary aggregation.

HAP1 uses 164 top-10 queries, SPIDR 449 top-10 gene lists, Feng 13 top-5% queries and HuRI 2,111 top-10 queries with at least one HuRI-positive partner. For HuRI, per-list behavior is computed before labels are loaded on all 2,424 archived lists; the predefined positive-containing subset is selected only for the final primary aggregation. The all-2,424-list closure is 0.573 [0.559, 0.585]. Thus the score prediction and per-list overlap do not use target hit values, while the primary HuRI reporting population is label-defined.

**Table S14:** Primary LR behavior prediction. Overlaps and closures are averaged over ten redraws. Intervals are two-stage 95% percentile intervals. Sources: `E11_behavior.csv`.

| Pool | Lists | $O(R, M)$ | $O(\widehat{M}, M)$ | Closure | 95% interval |
|---|---|---|---|---|---|
| HAP1 | 164 | 0.179 | 0.665 | 0.592 | [0.560, 0.625] |
| SPIDR | 449 | 0.356 | 0.777 | 0.654 | [0.640, 0.669] |
| Feng | 13 | 0.436 | 0.747 | 0.551 | [0.472, 0.617] |
| HuRI | 2,111 | 0.327 | 0.711 | 0.571 | [0.558, 0.585] |

**Retrieval-change endpoint (P2).** After behavior predictions and list overlaps are stored, the existing labels are used to calculate $\widehat{\Delta} = H(\widehat{s}_M) - H(s_R)$ and $\Delta = H(s_M) - H(s_R)$, where $H$ is the sum of tie-weighted hits across the pool's query lists. The criterion requires matching signs in all four pools and a predicted-change 95% interval excluding zero in each. P1 and P2 share 2,000 paired two-stage bootstrap samples (seed 20261024): redraw indices and query lists are resampled, with lists stratified by CV3 fold except for the 13-query Feng panel. These are conditional intervals, not independent biological-screen replications or a project-wide multiplicity correction.

**Table S15:** LR retrieval-change prediction. Values are average changes in hit incidences relative to Random training, with 95% intervals. Counts have different budgets and numbers of query lists across pools. Sources: `E11_retrieval_contrasts.csv`.

| Pool | Retrained $\Delta$ [interval] | Predicted $\widehat{\Delta}$ [interval] | $\widehat{\Delta}/\Delta$ |
|---|---|---|---|
| HAP1 | 12.2 [5.4, 20.8] | 11.1 [4.4, 19.6] | 0.91 |
| SPIDR | −32.5 [−60.6, −3.7] | −18.9 [−45.6, 8.5] | 0.58 |
| Feng | −8.2 [−14.6, −2.3] | −8.6 [−16.3, −1.3] | 1.05 |
| HuRI | 61.5 [40.6, 83.0] | 56.7 [37.4, 76.4] | 0.92 |

P1 passes its declared criterion (mean closure 0.592). P2 does not: all point-estimate directions agree, but the SPIDR predicted-change interval crosses zero. The paired intervals for $\Delta - \widehat{\Delta}$ include zero in all four pools; this is not proof of equivalence or acceptable prediction error. Descriptive SCHEMATIC top-3 retrieval over 149 gene lists has closure 0.581 [0.533, 0.629], but both true and predicted retrieval-change intervals cross zero.

**Scope across features and learners.** For SL/PPI, median standardized cosine similarity between the discriminator coefficient vector and $\beta_R - \beta_M$ is 0.983/0.911 overall and 0.993/0.970 in the endpoint block. Endpoint-only correction gives closure 0.420, 0.506, 0.344 and 0.474 across HAP1, SPIDR, Feng and HuRI; pair-only correction gives 0.005, −0.050, −0.545 and 0.143. One-dimensional correction gives 0.188, 0.185, 0.003 and 0.297, and the endpoint-ablated model 0.111, 0.075, 0.058 and 0.031. These comparisons support a multidimensional negative-distribution prediction, rather than merely reproducing one degree scalar.

With logistic density correction, HGB closures are 0.096, 0.080, 0.003 and 0.167; a nonlinear HGB discriminator changes them to 0.119, 0.116, 0.011 and 0.190. MLP closures on the three SL panels are 0.075, 0.092 and −0.003. HGB's predicted retrieval changes have the correct sign and intervals excluding zero in all four pools, despite low list closure. These secondary results separate coarse retrieval direction from accurate identification of the changed list; they do not extend the primary LR behavior claim to all learners.

The exploratory SLMGAE comparison uses validation-selected states (three initializations, one fixed negative set per arm). Because reconstruction scores are not logits and some are negative, its separately frozen score transformation is $\log(\max(s_R, 0) + 10^{-6}) - \mathrm{clip}(\log \widehat{r})$. Original-score rankings define the true lists. Top-10 closures are −1.846 [−2.271, −1.457] in HAP1 and −2.968 [−3.399, −2.643] in SPIDR. SPIDR's predicted change is −133.7 hits [−171.7, −96.3], versus −32.0 observed. This transformation fails to predict SLMGAE list behavior and is not presented as a general post-processing repair. The input views, score semantics and initialization-only replication differ from the primary LR setting.

### S16. Predictions frozen before training the matched model

**New relation tasks and sequence.** E12a uses two knowledge-graph relations not previously used as outcome labels in this project: evolutionary-rate covariance (COVARIES_GcG) and a LINCS-derived regulation relation (REGULATES_GrG). Self-loops are removed and relations are represented as unique unordered pairs, including symmetrization of REGULATES. COVARIES contains 16,972 pairs among 5,357 genes and REGULATES 146,542 among 5,803 genes. The same 9,845-gene CV3 partition defines source-positive pairs and held-out pairs. Each relation's five directly encoding features (edge indicator, common-neighbour count, neighbour Jaccard and two endpoint-degree features) are removed, leaving 54. Remaining features share the knowledge-graph source and may retain indirect relation information.

Each task uses ten negative redraws, equal positive and negative counts, and both LR and HGB with the settings above. Random/matched sampling seeds are $12001 + r/13001 + r$ for COVARIES and $14001 + r/15001 + r$ for REGULATES ($r = 0, \ldots, 9$). Matching uses the source-training graph's ordered endpoint-degree bins and excludes all known positives for that relation. The discriminator and clipping specification is the same as in S15. The source-only discriminator AUROC gate requires a mean of at least 0.55; observed means are 0.597 and 0.906. COVARIES requires 90 adjacent-bin fallback pairs among 543,350 matched negatives across all fits; REGULATES requires none.

Stage A fits Random models and negative-distribution discriminators, stores corrected predictions and list summaries, and freezes their hashes. Stage B verifies those hashes before fitting any matched model and evaluates the unchanged predictions once. The prediction freeze is dated 2026-10-28 06:03:29 UTC; Stage B starts at 06:04:19 UTC. The held-out evaluation universe contains all 1,969 test genes per fold, each ranking the other 1,968 genes. All 9,845 genes are queries, without selection using held-out labels. The primary endpoint is the S15 list-closure statistic with top 10 and 2,000 paired bootstrap samples (seed 20261024). Passing requires closure at least 0.5 and a lower 95% interval bound of at least 0.4. COVARIES is the primary task; REGULATES has the same criterion but is reported separately. This is prediction-before-fitting in a computational test, not prospective biological validation.

**Table S16:** Frozen predictions on both new relation tasks. All comparisons use 9,845 queries and ten redraws. HGB is secondary. Sources: `E12A_results.csv`.

| Relation | Learner | $O(R, M)$ | $O(\widehat{M}, M)$ | Closure [95% interval] | Criterion |
|---|---|---|---|---|---|
| COVARIES | LR | 0.252 | 0.767 | 0.689 [0.678, 0.699] | Met |
| COVARIES | HGB | 0.060 | 0.062 | 0.002 [−0.001, 0.004] | Not met |
| REGULATES | LR | 0.405 | 0.564 | 0.267 [0.260, 0.274] | Not met |
| REGULATES | HGB | 0.160 | 0.098 | −0.074 [−0.082, −0.066] | Not met |

Coefficient-direction agreement alone does not explain the ranking outcomes. For LR, standardized coefficient cosines are 0.965 (COVARIES) and 0.994 (REGULATES), while norm ratios are 0.851 and 0.938. REGULATES' score–endpoint coupling changes from 0.787 to 0.261 after true retraining, but the prediction moves it to 0.021. COVARIES coupling changes from −0.147 to −0.135, whereas the prediction is −0.161; the true change interval crosses zero. High coefficient alignment therefore does not guarantee precise behavior in the extreme ranking tail.

Held-out retrieval uses each relation's own test positives, not an independent experiment. For COVARIES LR, the retrained hit change is 1.0 [−13.9, 15.7] and the predicted change 3.0 [−11.6, 17.8]; list prediction success does not establish a retrieval gain. For REGULATES LR, the true change is −791.0 [−914.7, −662.2] and the predicted change −600.7 [−720.3, −482.1]. Corresponding HGB changes are −7.4 [−22.4, 6.2] versus predicted −8.2 [−16.7, −0.2] for COVARIES, and −1785.4 [−1928.5, −1644.8] versus predicted −2166.8 [−2306.7, −2035.5] for REGULATES. All are query-ranking incidences, not unique pair discoveries. The primary task supports the specified LR behavior forecast, while the second task and HGB results delimit its transfer. Computational folds and redraws do not remove shared-source dependence.

### S17. Density-ratio interpretation, simulation and decision-rule limits

**Population identity and feature projection.** With common positive distribution $P_+(x)$, negative distribution $Q(x)$ and equal class priors, the population-optimal classification logit is

$$f_Q^*(x) = \log \frac{P_+(x)}{Q(x)}.$$

This density-ratio interpretation is established, including in noise-contrastive estimation [13]; it is not a new theorem claimed by this study. For Random negatives $U$ and matched negatives $M$, wherever the relevant densities are positive,

$$f_M^*(x) = f_U^*(x) - \log \frac{M(x)}{U(x)}, \qquad f_{Q_p}^*(x) = f_U^*(x) - \log[(1-p) + p\, r(x)],$$

where $Q_p = (1-p)U + pM$ and $r = M/U$. Class-prior differences would add an intercept. Matching on source-graph bins and learning from auxiliary features are different operations: the relevant ratio is the distribution induced in the learner's feature space. Source-only density estimation can therefore be informative about held-out genes even when their degrees in the source training graph are all zero. Exact population logits are not guaranteed for finite, regularized or misspecified learners. E11 and E12a test the resulting approximation empirically rather than assuming it.

In the special factorization $M(e, z) = P_+(e)U(z \mid e)$ for endpoint attributes $e$ and pair attributes $z$, the logit difference is $h(e) = \log[P_+(e)/U(e)]$. Relative to the fully matched logit, mixed negatives leave the residual term

$$h_p(e) = -\log\{(1-p)e^{-h(e)} + p\}.$$

It approaches zero as $p$ approaches one and, for $0 < p < 1$, has derivative with respect to $h$ between zero and one. This statement about the endpoint term does not imply that every rank correlation between the *total* score and degree must decrease monotonically: the pair term and its dependence on endpoint features also matter. Empirical coupling curves are reported as observations, not universal consequences of this identity.

For a benchmark with positives $P_+$ and test negatives $T$, likelihood-ratio ordering $P_+/T$ maximizes discrimination at a fixed false-positive rate in the ideal population setting. Thus models attaining $f_U^*$

and $f_M^*$ would be optimal on tests using their respective negative distributions. This explains why test composition can favor different score orderings, but does not establish that finite fitted models will necessarily achieve or reverse those optima. Experimental retrieval has a different candidate distribution and target label process.

**Simulation and a conditional threshold.** The archived simulation uses 2,000 genes partitioned into five folds. Each gene has a study-bias attribute and a four-dimensional latent vector; latent pair affinity is half their inner product. Approximately 6,000 source positives are sampled with log odds depending on affinity plus the two genes' study attributes. Observed endpoint and pair features are noisy measurements of these quantities. Each test fold supplies 40 queries with roughly 400 candidate partners, with experimental hit prevalence targeted at 1%. A parameter $\gamma$ changes the target label's dependence on partner study attribute. Fixed LR models compare Random and source-training-degree-matched negatives at top 10. Eleven $\gamma$ values and 12 repeats yield 132 runs; the simulation is a constructed mechanism illustration, not an independent biological validation.

Under an additional local Gaussian approximation, write target latent signal as $bf_M + ch + \epsilon$, assume $f_M$ and $h$ independent, and set $\beta = b\sigma_f > 0$, $\Gamma = c\sigma_h$ and $\rho = \sigma_h/\sigma_f > 0$. If retrieval increases with the score's correlation to that latent signal, ranking by $f_M + h$ rather than $f_M$ is favored when

$$\frac{\Gamma}{\beta} > \kappa(\rho) = \frac{\sqrt{1+\rho^2}-1}{\rho}.$$

This conditional result illustrates why a positive degree-only gain does not suffice to select Random training: its contribution can be positive but too weak for the source-trained weight. Replacing the latent-signal ratio by $G_{\text{degree}}/G_M$ gives a heuristic decision rule, not an exact identity. It requires the target labels needed to calculate both gains and is not a label-free prescription for a new screen. A correlation-based estimate of $\rho$ additionally relies on the Gaussian independence assumptions; empirical rank coupling is only a proxy.

**Table S17:** Archived synthetic-data comparison. $\Delta$ is matched-minus-Random top-10 precision gain in pp (mean $\pm$ SD over 12 repetitions). The last two columns are the fractions of simulated repetitions with correctly predicted effect direction. These are simulation accuracies, not clinical or biological prediction accuracies. Source: `C_sim_summary.csv`.

| $\gamma$ | $\Delta$ (pp) | Sign rule | Threshold heuristic |
|---:|---:|---:|---:|
| $-1.00$ | $3.00 \pm 0.39$ | 1.00 | 1.00 |
| $-0.50$ | $2.68 \pm 0.31$ | 1.00 | 1.00 |
| $-0.25$ | $2.11 \pm 0.45$ | 0.75 | 1.00 |
| 0.00 | $1.64 \pm 0.45$ | 0.08 | 1.00 |
| 0.10 | $1.29 \pm 0.53$ | 0.00 | 1.00 |
| 0.25 | $0.64 \pm 0.38$ | 0.00 | 0.92 |
| 0.40 | $0.17 \pm 0.54$ | 0.42 | 0.58 |
| 0.60 | $-0.75 \pm 0.33$ | 1.00 | 1.00 |
| 0.80 | $-2.20 \pm 0.71$ | 1.00 | 1.00 |
| 1.00 | $-3.38 \pm 0.72$ | 1.00 | 1.00 |
| 1.50 | $-6.74 \pm 0.70$ | 1.00 | 1.00 |

**Retrospective empirical check.** The corrected source table combines 51 previously examined screen–budget–learner cells. The sign rule agrees with 41 point-estimate directions and the threshold heuristic with 45. The four additional agreements concern HAP1 and SCHEMATIC at a 10% budget for LR/HGB; all four effect intervals include zero. In the 20 corrected cells whose effect intervals exclude zero, both rules agree with all 20 directions. These cells share pools, labels and fitted models, so the counts are neither independent validations nor a demonstrated improvement in decision accuracy. The corrected Feng mapping changes cell-level values, but not the aggregate 41/51 and 45/51 counts; the corrected table is `C_kappa_posthoc_amend1.csv`.

The HuRI result in Section S13 already shows that a degree-only sign rule can fail even when endpoint-coefficient effects transfer. Neither the illustrative threshold nor retrospective cell counts establish a universal replacement rule. The role of this theoretical material is to motivate the negative-distribution predictor, identify assumptions and explain why predicting model behavior remains distinct from predicting which training design maximizes experimental retrieval.
